# The human disease gene *CLEC16A* encodes an intrinsically disordered protein region required for mitochondrial quality control

**DOI:** 10.1101/2021.09.03.458272

**Authors:** Morgan A. Gingerich, Xueying Liu, Biaoxin Chai, Gemma L. Pearson, Michael P. Vincent, Tracy Stromer, Jie Zhu, Vaibhav Sidarala, Aaron Renberg, Debashish Sahu, Daniel J. Klionsky, Santiago Schnell, Scott A. Soleimanpour

**Affiliations:** Department of Internal Medicine and Division of Metabolism, Endocrinology & Diabetes, University of Michigan, Ann Arbor, MI; Program in Cellular and Molecular Biology, University of Michigan, Ann Arbor, MI; Institute of Metabolism and Endocrinology, The Second Xiangya Hospital, Central South University, Changsha, China; Department of Molecular and Integrative Physiology, Department of Computational Medicine and Bioinformatics, University of Michigan, Ann Arbor, MI; University of Michigan BioNMR Core Facility, Ann Arbor, MI; Life Sciences Institute and Department of Molecular, Cellular, and Developmental Biology, University of Michigan, Ann Arbor, MI; VA Ann Arbor Health Care System, Ann Arbor, MI

**Author notes:** **Corresponding Author** Scott A. Soleimanpour, MD, 1000 Wall Street, Brehm Tower Room 5317, Ann Arbor, MI 48105.

## Abstract

CLEC16A regulates mitochondrial health through mitophagy and is associated with over 20 human diseases. While CLEC16A has ubiquitin ligase activity, the key structural and functional regions of CLEC16A, and their relevance for human disease, remain unknown. Here, we report that a disease-associated CLEC16A variant lacks a C-terminal intrinsically disordered protein region (IDPR) that is critical for mitochondrial quality control. Using carbon detect NMR, we find that the CLEC16A C terminus lacks secondary structure, validating the presence of an IDPR. Loss of the CLEC16A C-terminal IDPR *in vivo* impairs pancreatic β-cell mitophagy, mitochondrial function, and glucose-stimulated insulin secretion, ultimately causing glucose intolerance. Deletion of the CLEC16A C-terminal IDPR increases its self-ubiquitination and destabilizes CLEC16A, thus impairing formation of a critical CLEC16A-dependent mitophagy complex. Importantly, CLEC16A stability is dependent on proline bias within the C-terminal IDPR, but not amino acid sequence order or charge. Together, we clarify how an IDPR in CLEC16A prevents diabetes, thus implicating the disruption of IDPRs as novel pathological contributors to diabetes and other CLEC16A-associated diseases.

## INTRODUCTION

*CLEC16A* (*C-type lectin domain containing 16A*) is a gene associated with nearly 20 human diseases, including type 1 diabetes, cardiovascular disease, and multiple sclerosis (1-5). *CLEC16A* encodes an E3 ubiquitin ligase which regulates mitochondrial quality control by clearing damaged or aged mitochondria through a type of selective autophagy, termed mitophagy (6; 7). CLEC16A forms and stabilizes a tripartite mitophagy complex with the E3 ubiquitin ligase RNF41/Nrdp1 and the deubiquitinase USP8, which together regulate mitophagic flux by controlling the activity of the mitophagy-effector Parkin (PRKN; (8; 9)). Genetic or pharmacologic disruption of CLEC16A impairs mitochondrial function and insulin secretion in pancreatic β-cells, leading to hyperglycemia and diabetes (6; 8-11). Whereas CLEC16A has a well-defined role in regulating mitophagy, the critical structural and functional regions of CLEC16A, and their relevance to disease pathogenesis, are unknown.

In humans, *CLEC16A* encodes two consensus isoforms generated by alternative splicing, a 24-exon full-length transcript, and a shorter 21-exon transcript variant that encodes a protein with a truncated C terminus. A single nucleotide polymorphism (SNP) that is associated with increased expression of the shorter, C-terminal truncated CLEC16A isoform in the thymus is known to increase risk for both diabetes and multiple sclerosis (12; 13). Additionally, several *CLEC16A* disease-associated SNPs promote increased expression of the C-terminal deficient isoform in a variety of cell types, acting as splicing quantitative trait loci (14; 15). These observations led us to hypothesize that the CLEC16A C terminus directly contributes to CLEC16A function and protection from disease.

The C terminus of CLEC16A does not share homology to known protein domains; rather, it is suggested to be an intrinsically disordered protein region (IDPR). IDPRs lack secondary structure and exist as an ensemble of flexible inter-converting conformations (16; 17). Although the roles of IDPRs have only recently begun to be explored, these domains support critical biological functions, including signal transduction, protein complex assembly, and protein stability (16-20). Disruption or dysregulation of IDPRs has been associated with several human diseases, further suggesting their functional importance (21-25). However, the molecular mechanisms connecting loss of IDPRs to disease pathogenesis are poorly understood. Moreover, the genetic, functional, and physiological connections between disruption to IDPRs and diabetes pathogenesis are unknown.

Here, we investigate how the putative CLEC16A C-terminal IDPR contributes to CLEC16A structure, function, and glucose homeostasis. We validate that the C-terminal region lost in the *CLEC16A* disease variant is an IDPR using rigorous *in silico*, biochemical, and NMR-based approaches. The CLEC16A C-terminal IDPR is crucial for glucose homeostasis by promoting glucose-stimulated insulin secretion, mitochondrial function, and mitophagy in β-cells. Loss of the CLEC16A C-terminal IDPR reduces CLEC16A stability due to increased self-ubiquitination and degradation, which impairs assembly of the CLEC16A-Nrdp1-USP8 tripartite mitophagy complex. Finally, we determine that the CLEC16A C-terminal IDPR depends on its proline enrichment, but not primary amino acid sequence order or charge, to stabilize CLEC16A. Together, we define the molecular mechanisms by which a disrupted IDPR in a *CLEC16A* disease variant destabilizes CLEC16A, impairs β-cell mitophagy, and contributes to diabetes.

## RESULTS

### The CLEC16A C terminus is an intrinsically disordered protein region

To understand the importance of the C-terminal region of CLEC16A that is lost in the disease variant, we first investigated the presence of conserved functional domains. Bioinformatic software was used to predict domains against the Pfam database of known functional protein domains (26). This analysis did not identify any conserved functional domains in the CLEC16A C terminus (Figure 1A) (26). Interestingly, the CLEC16A C terminus was predicted to be an intrinsically disordered protein region (IDPR; Figure 1A). CLEC16A also contains an N-terminal “FPL” domain (Figure 1A). However, the 150 amino acid “FPL” domain has no known function and is only found in CLEC16A orthologs. To further evaluate the presence of a putative IDPR within the CLEC16A C terminus, we calculated the mean disorder score from three independent disorder-prediction algorithms IUPRED2, Disprot VSL2B, and DISOPRED 3.1 (27-29). This analysis identified two conserved predicted IDPRs within human and mouse CLEC16A, including the C-terminal region that is lost in the CLEC16A disease variant (Figures 1B and S1). As a complementary approach, we compared the amino acid composition of the mouse CLEC16A C terminus to that of validated IDPRs. Proline and serine are well-known disorder-promoting amino acids and are significantly enriched in the disordered protein database Disprot relative to the expected distribution of amino acids found in nature, approximated by the SwissProt database (Figure S2) (30-32). The CLEC16A C terminus is significantly enriched in the disorder-promoting amino acids proline and serine relative to the SwissProt Database (Figures 1C and D). Thus, several independent approaches predict the CLEC16A C terminus to be an IDPR.

**Figure 1.**
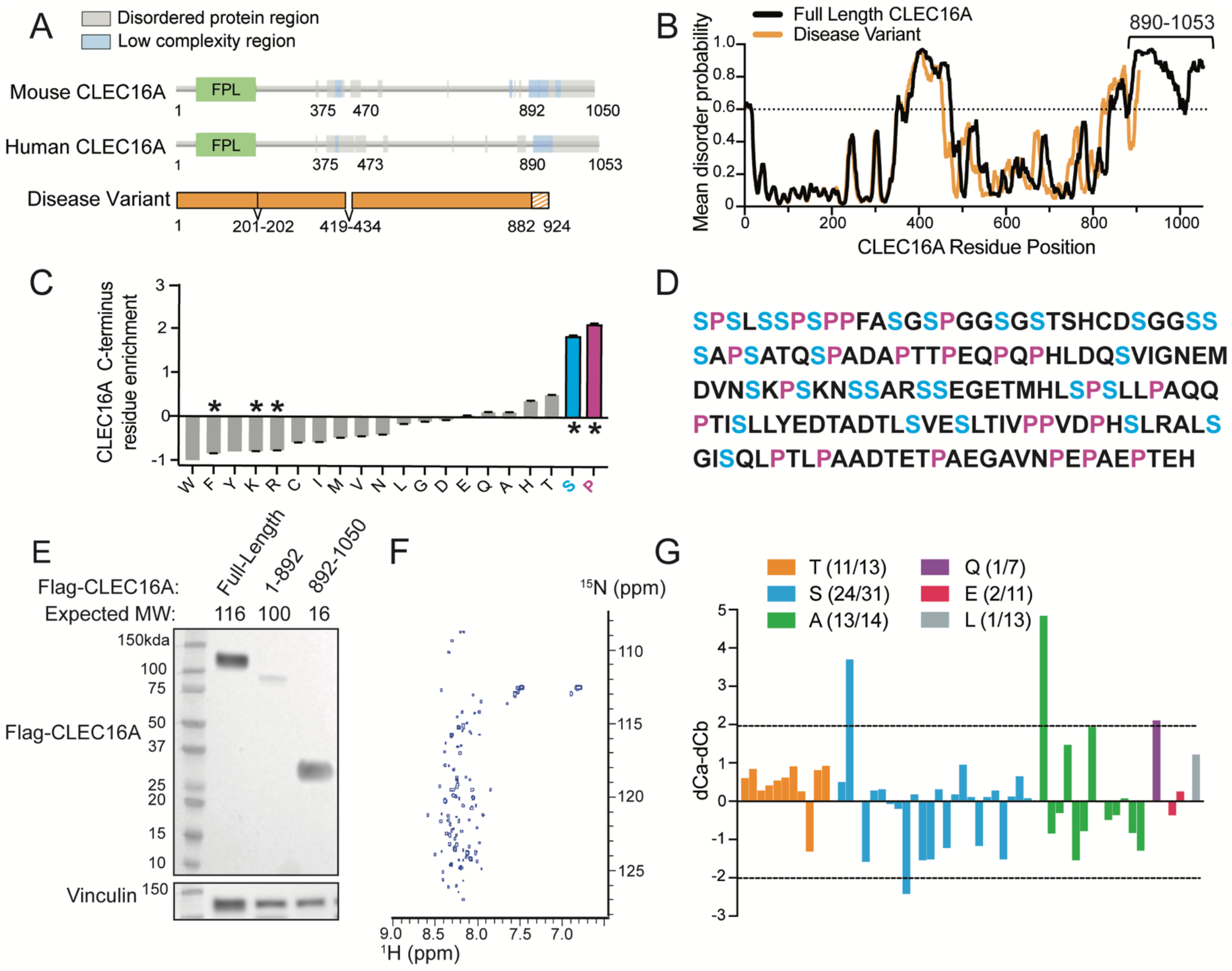
A human CLEC16A disease variant lacks a C-terminal IDPR. **(A)** Domain prediction of the human CLEC16A disease variant and human full-length CLEC16A generated by Pfam. Human CLEC16A disease variant is aligned beneath, shaded region represents sequence differing from human full-length CLEC16A. ‘FPL’ is a domain of unknown function enriched in the amino acids phenylalanine (F), proline (P), and leucine (L). **(B)** Mean disorder score from IUPred 2, Disprot VSL2B, and DISOPRED 3.1 of full-length CLEC16A and the human CLEC16A disease variant. Putative disordered regions were identified with a probability threshold of >0.6. **(C)** Residue composition bias of the mouse CLEC16A C-terminus generated with Composition Profiler, comparing residue enrichment of the CLEC16A C-terminus vs SWISS-PROT 51 database. Significantly enriched serine (blue) and proline (pink) residues are highlighted. * p<0.05. **(D)** Mouse CLEC16A C terminus residues (AA 892-1050) are listed in FASTA format. Serine residues highlighted in blue, and proline residues are in pink. **(E)** Representative western blot (WB) in HEK293T cells transfected with Flag-CLEC16A constructs (full length CLEC16A, CLEC16A lacking the C terminus (CLEC16A ΔC, AA 1-892), or CLEC16A C terminus only (AA 892-1050)). Vinculin serves as a loading control. n=3/group. **(F)** ^1^H-^15^N HSQC spectra of recombinant CLEC16A C terminus (AA 892-1050). Spectra clusters near ^1^H 8ppm, consistent with being an IDPR. **(G)** Assessment of secondary structure propensity of residue types identified by NMR. Secondary structure is shown as the difference between ^13^C_α_ and ^13^C_β_ secondary chemical shifts. Stretches of values > 2ppm indicate α-helix, values < 2ppm indicate β-sheet.

To validate the presence of an IDPR within the CLEC16A C terminus, we used both biochemical and biophysical techniques. IDPRs commonly exist in an extended conformation and are often enriched in charged residues, resulting in poor binding to sodium dodecyl sulfate (SDS) and slower migration on an SDS-polyacrylamide gel (33; 34). While Flag epitope-tagged full-length CLEC16A and a C-terminal deficient CLEC16A mutant (CLEC16A ΔC) migrated at their expected molecular weights, a CLEC16A construct encoding only the C terminus migrated more slowly, at nearly double its expected molecular weight (Figure 1E). We next investigated CLEC16A C-terminal structure using nuclear magnetic resonance (NMR). In an IDPR, backbone hydrogen atoms do not participate in hydrogen bonds that generate secondary structure, and instead are found in a flexible extended conformation. Thus, all backbone hydrogen atoms in an IDPR are in a similar chemical environment and are tightly clustered in the hydrogen dimension of an ^1^H-^15^N heteronuclear single quantum coherence (HSQC)-NMR spectrum (35). The HSQC-NMR spectrum of the CLEC16A C terminus was tightly clustered in the hydrogen dimension between 8-8.5 ppm, which strongly suggests that the CLEC16A C terminus is an IDPR (Figure 1F).

To complement our macroscopic view of the CLEC16A C terminus as an IDPR, we investigated structural features at the resolution of single amino acids using carbon-detect NMR. While we were unable to assign NMR resonance peaks to specific amino acids in the CLEC16A C terminus due to its repetitive sequence and complex spectrum, we instead were able to use carbon-detect amino-acid specific (CAS) experiments to assign amino acid types to NMR resonance peaks (35). Using CAS, we assigned amino acid types to 52 resonance peaks within the putative CLEC16A C-terminal IDPR, with identified residue types distributed throughout the C terminus (Figures 1D and G). The secondary structure character of assigned resonance peaks was measured as the difference between the offset of the chemical shift of ^13^C-α and ^13^C-β nuclei relative to that expected for each amino acid type in an IDPR (Figure 1G). Using this approach, stretches of values greater than or less than 2ppm suggest α-helical or β-sheet character, respectively (35; 36). Within the CLEC16A C-terminal fragment, only 4 of the 52 identified residues had character consistent with secondary structure, indicating that this region is highly disordered (Figure 1G). Together, our *in silico*, biochemical, and biophysical studies strongly support the hypothesis that the C-terminal region lost in the CLEC16A disease variant is an IDPR.

### The Clec16a C-terminal IDPR is required for glucose homeostasis and β-cell function

After confirming that the C terminus of CLEC16A disrupted in the human disease variant is an IDPR, we questioned whether this region impacts the role of CLEC16A in glucose homeostasis and β-cell function. To study the function of the CLEC16A C terminus *in vivo*, we used a genetic mouse model lacking the CLEC16A C terminus. The *Clec16a* curly tail mutant (*Clec16a*^curt^) mouse carries a spontaneous four base-pair deletion within exon 22, leading to a frameshift mutation that is followed by alternatively translated residues and a premature stop codon (Figures 2A-C) (37-39). The *Clec16a*^curt^ mutant encodes a truncated protein lacking the C-terminal IDPR, which is highly similar to the CLEC16A disease variant (Figures 2A and S1). The truncated C-terminal deficient CLEC16A^curt^ protein was detectable in mouse embryonic fibroblasts (MEFs) and pancreatic islets, albeit at lower levels than full length wild type (WT) CLEC16A (Figures 2D,E). Additionally, antisera specifically recognizing the C terminus of CLEC16A were unable to detect the CLEC16A^curt^ mutant protein (Figure 2F). *Clec16a*^curt/curt^ mice were born at a normal Mendelian ratio and had reduced body weight, motor impairments, and early lethality around 6 months of age, similar to previous reports (data not shown; (37-39)). *Clec16a*^curt/+^ heterozygous mice were indistinguishable from wild type littermates (data not shown).

**Figure 2.**
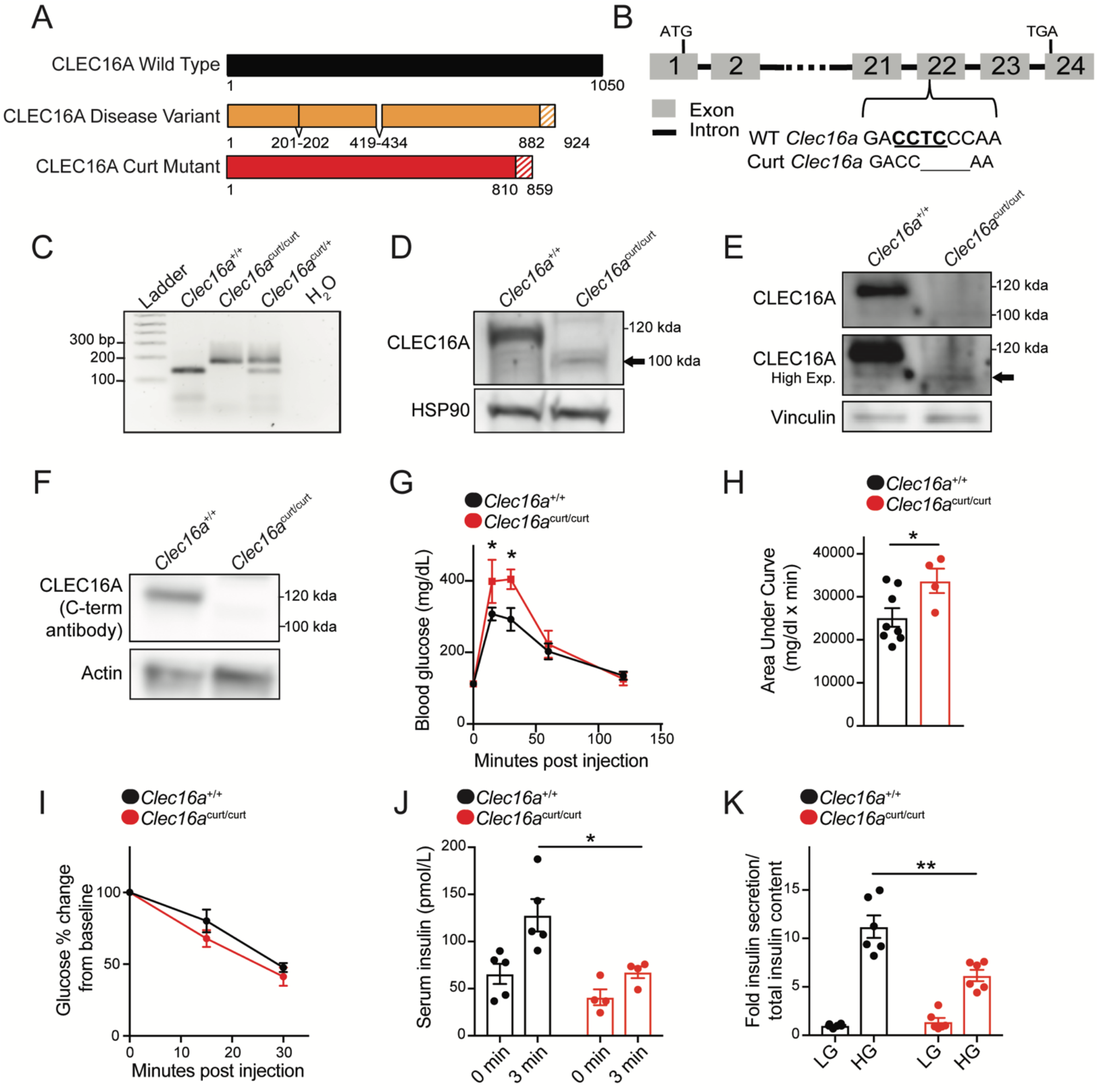
The CLEC16A C-terminus maintains glucose homeostasis and β-cell function. **(A)** Schematic of the CLEC16A^curt^ protein and human CLEC16A disease variant. Shaded region represents sequence differing from full-length mouse CLEC16A. **(B)** Schematic of *Clec16a* locus, highlighting the 4bp deletion within exon 22 of *Clec16a*^curt^ mice. The *MnlI* recognition sequence is underlined. **(C)** Representative image of *MnlI* restriction digest following PCR amplification of genomic DNA containing *Clec16a* exon 22 as visualized by agarose gel electrophoresis. The *Clec16a*^curt^ mutation eliminates the *MnlI* digestion site. n=4/group. **(D)** Representative image of CLEC16A protein levels determined by WB in *Clec16a*^+/+^ and *Clec16a*^curt/curt^ MEFs utilizing CLEC16A anti-sera raised against an internal CLEC16A peptide. Hsp90 serves as loading control. n=3/group. **(E)** Representative image of CLEC16A protein levels in *Clec16a*^+/+^ and *Clec16a*^curt/curt^ isolated islets utilizing CLEC16A anti-sera raised against an internal CLEC16A peptide. n=3/group. **(F)** Representative image of CLEC16A protein levels determined by WB in *Clec16a*^+/+^ and *Clec16a*^curt/curt^ MEFs utilizing CLEC16A anti-sera raised against a C-terminal CLEC16A peptide. Beta actin serves as loading control. n=3/group. **(G)** Blood glucose concentrations measured during an intraperitoneal glucose tolerance test (IPGTT) of 16-week-old *Clec16a*^+/+^ and *Clec16a*^curt/curt^ littermates. n=8 *Clec16a*^+/+^, n=4 *Clec16a*^curt/curt^. **(H)** Area under curve from IPGTT in Figure 2G. **(I)** Blood glucose concentrations during insulin tolerance test of 9-week-old *Clec16a*^+/+^ and *Clec16a*^curt/curt^ littermates. n=3-4/group. **(J)** Serum insulin measured during *in vivo* glucose-stimulated insulin release in 11-week-old *Clec16a*^+/+^ and *Clec16a*^curt/curt^ littermates. n=4-5/group. **(K)** Fold glucose-stimulated insulin secretion following static incubations in 1.67 mM and 16.7 mM glucose in isolated *Clec16a*^+/+^ and *Clec16a*^curt/curt^ islets from 11-week-old littermates. n=6/group. *p<0.05 **p<0.01.

To investigate the impact of the CLEC16A C terminus on glucose homeostasis, we first performed an intraperitoneal glucose tolerance test (IPGTT). *Clec16a*^curt/curt^ mice were glucose intolerant compared to littermate controls (Figures 2G,H). Impaired glucose tolerance can result from reduced sensitivity of peripheral tissues to insulin or reduced insulin secretion in response to glucose. *Clec16a*^curt/curt^ mice had no reduction in insulin sensitivity during an insulin tolerance test (Figure 2I). However, *Clec16a*^curt/curt^ mice secreted significantly less insulin in response to glucose stimulation both *in vivo* and in isolated islets (Figures 2J,K), which indicated that glucose intolerance was due to reduced β-cell insulin secretion. When compared to previous studies on β-cell or pancreas-specific *Clec16a* knockout mice, *Clec16a*^curt/curt^ mice had a similar, yet more modest phenotype of impaired glucose homeostasis and insulin secretion (6; 8). These results suggest that the CLEC16A^curt^ truncated protein was less functional than full-length WT CLEC16A, indicating that the CLEC16A C-terminal IDPR is critical for optimal β-cell function.

We next evaluated whether reduced insulin secretion in *Clec16a*^*c*urt/curt^ mice was due to reduced β-cell mass or insulin content. *Clec16a*^curt/curt^ mice had no significant changes in β-cell mass or pancreatic insulin content proportional to body weight or pancreas weight, respectively, despite reduced body weight in *Clec16a*^curt/curt^ mutants (Figure S3). Together, these studies indicate that the CLEC16A C-terminal IDPR regulates glucose homeostasis through control of β-cell insulin secretion.

### The CLEC16A C terminus regulates β-cell mitophagy

CLEC16A-mediated mitophagy maintains mitochondrial health and function in β cells, which is essential to fuel glucose-stimulated insulin secretion (6; 40). Thus, we next assessed how loss of the CLEC16A C-terminal IDPR impacted β-cell mitochondria. *Clec16a*^curt/curt^ β-cells displayed dysmorphic mitochondria with disorganized cristae when compared to littermate controls, as visualized by transmission electron microscopy (TEM) (Figure 3A). Consistent with these morphological defects, mitochondrial function was reduced in isolated *Clec16a*^curt/curt^ islets, measured by glucose-stimulated oxygen consumption rates (Figure 3B). *Clec16a*^curt/curt^ islets also had reduced FCCP-stimulated maximal oxygen consumption rates, while displaying no differences in glycolysis as measured by extracellular acidification rates, when compared to littermate controls (data not shown).

**Figure 3.**
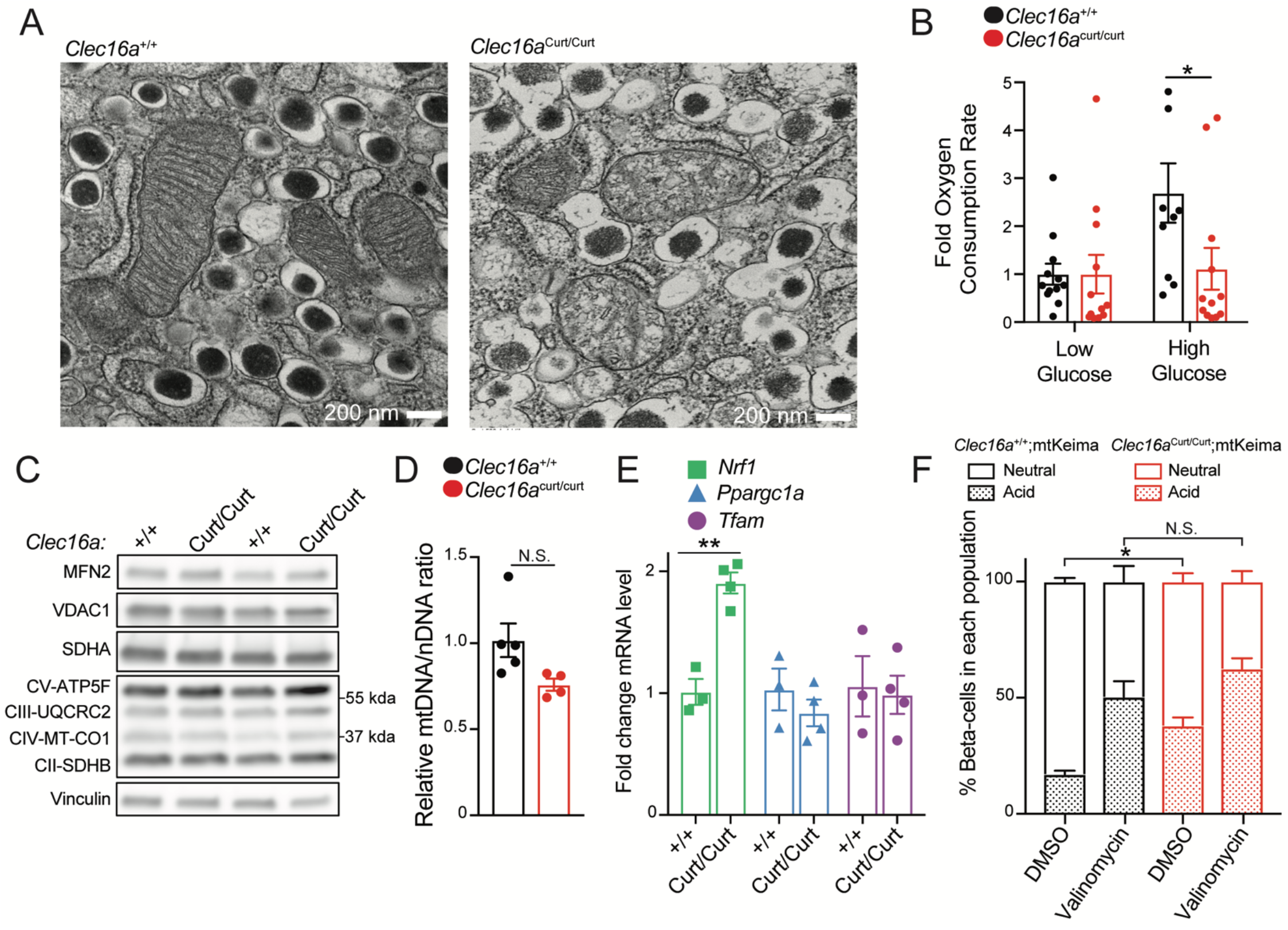
The CLEC16A C-terminus maintains β-cell mitochondrial function, morphology, and turnover. **(A)** Representative transmission electron microscopy (TEM) images of β-cells in isolated islets from 11-week-old *Clec16a*^+/+^ and *Clec16a*^curt/curt^ islets. *Clec16a* ^*curt/curt*^ islets have distorted mitochondria cristae. n=4/group. **(B)** Relative oxygen consumption rate (OCR) in isolated *Clec16a*^+/+^ and *Clec16a*^curt/curt^ islets measured after exposure to low glucose (1.67 mM) and high glucose (16.7 mM). n=12/group. **(C)** Representative WB of select mitochondrial proteins from isolated *Clec16a*^+/+^ and *Clec16a*^curt/curt^ islets. n=4/group. **(D)** Relative mitochondrial DNA (mtDNA) to nuclear DNA (nDNA) ratio in isolated islets from *Clec16a*^+/+^ and *Clec16a*^curt/curt^ mice. n=4-5/group. **(E)** Relative mRNA levels of mitochondrial biogenesis markers (normalized to *Hprt* expression) in isolated islets from 12-week old *Clec16a*^+/+^ and *Clec16a*^curt/curt^ mice. n=3-4/group. **(F)** Flow cytometric analysis of β-cells from dissociated islets of 12-week old *Clec16a*^+/+^;mtKeima and *Clec16a*^curt/curt^;mtKeima mice, indicating the relative distribution of β-cells with mitochondria in dominantly acidic or neutral compartments following exposure to 250 nM valinomycin or DMSO control for 3 hours. n=4/group. *p<0.05 **p<0.01.

Reduced mitochondrial function may stem from the presence of dysfunctional mitochondria or from reduced mitochondrial mass. To assess mitochondrial mass, we measured levels of key mitochondrial membrane proteins and electron transport chain subunits as well as mitochondrial DNA levels. *Clec16*a^curt/curt^ islets had no reductions in mitochondrial proteins or mitochondrial DNA content when compared to littermate controls (Figure 3C-D). We also measured expression of mitochondrial biogenesis markers *Ppargc1a/Pgc1*α and *Tfam* by qRT-PCR and found no significant differences in expression between groups (Figure 3E). Interestingly, expression of *Nrf1*, a mitochondrial biogenesis marker commonly induced following mitochondrial stress, was increased in *Clec16a*^curt/curt^ islets (41; 42). Collectively, these results demonstrate that *Clec16a*^curt/curt^ β-cells have reduced mitochondrial function that is not due to reduced mitochondrial mass, which could arise from impaired mitophagy.

To determine whether loss of the CLEC16A C-terminal IDPR affects β-cell mitophagy, we intercrossed the mt-Keima mitophagy reporter model with *Clec16a*^curt/curt^ mice. Mt-Keima mice express a fluorescently labeled pH-sensitive mitochondrial biosensor that shifts excitation spectra based on pH (43). Mt-Keima allows for detection of mitophagy as mitochondria are targeted to acidic autophagosomes/lysosomes for degradation. Flow cytometry of dissociated islets revealed that mt-Keima;*Clec16a*^curt/curt^ β-cells had an increase in cells with mitochondria in acidic compartments (Figure 3F). This suggests that *Clec16a*^*c*urt/curt^ β-cells accumulate mitochondria in acidic autophagosomes/lysosomes that may be incompletely cleared, consistent with previous observations following CLEC16A deficiency in β-cells (6). Additionally, membrane-engulfed mitochondria were observed in *Clec16a*^curt/curt^ β-cells by TEM, which were not found in littermate controls (Figure S4). Similar to *Clec16a*^curt/curt^ β-cells, mitochondria accumulated within acidic compartments in *Clec16a*^curt/curt^ MEFs, as measured using a cell permeable mitophagy reporter dye (Figure S5A-C). *Clec16a*^curt/curt^ MEFs also did not display differences in bulk autophagy markers including LC3 and p62, suggesting this defect is specific to mitophagy (Figure S5D). To determine if increased mitochondria within acidic compartments in *Clec16a*^curt/curt^ β-cells was due to impaired or enhanced mitophagy flux, we stimulated mitophagy using the mitochondrial ionophore valinomycin. Importantly, *Clec16a*^curt/curt^ β-cells did not demonstrate enhanced mitophagic flux following valinomycin treatment compared to *Clec16a*^*+/+*^ β-cells (Figure 3F). These results suggest the baseline accumulation of acidic mitochondria in *Clec16a*^curt/curt^ β-cells was due to impaired mitophagy. Therefore, the CLEC16A C-terminal IDPR maintains β-cell mitochondrial function and health through control of mitophagy.

### The CLEC16A C-terminal IDPR is required for CLEC16A stability and assembly of the tripartite mitophagy complex

Given the critical role of the CLEC16A C-terminal IDPR in regulating β-cell mitophagy, we questioned how this region impacts molecular functions of CLEC16A. IDPRs often promote protein turnover and destabilize proteins due to their enhanced accessibility for degradative post-translational modifications (18; 44; 45). However, both *Clec16a*^curt/curt^ MEFs and islets, which lack the CLEC16A C-terminal IDPR, had reduced levels of CLEC16A protein (Figures 2D,F). This led us to hypothesize that the C-terminal IDPR promotes CLEC16A stability, in contrast to the classic role of IDPRs in protein destabilization.

We investigated the role of the CLEC16A C terminus on protein stability using the previously described Flag-tagged CLEC16A ΔC construct which lacks the C-terminal IDPR (Figures 1E and 4A). CLEC16A ΔC had reduced protein levels relative to WT CLEC16A following transfection in HEK293T cells, despite no differences in mRNA expression (Figures 4B,C). To determine the contribution of the CLEC16A C terminus to the stability of the protein, we assessed CLEC16A protein turnover after inhibiting protein synthesis with cycloheximide. CLEC16A ΔC had accelerated turnover relative to wild-type CLEC16A (WT) following cycloheximide treatment, indicating that the C-terminal IDPR stabilizes CLEC16A (Figure 4D). Levels of WT CLEC16A and CLEC16A ΔC protein both increased when inhibiting the proteasome or lysosome with MG132 or bafilomycin A_1_, respectively, suggesting multiple pathways for CLEC16A clearance (Figure S6A).

**Figure 4.**
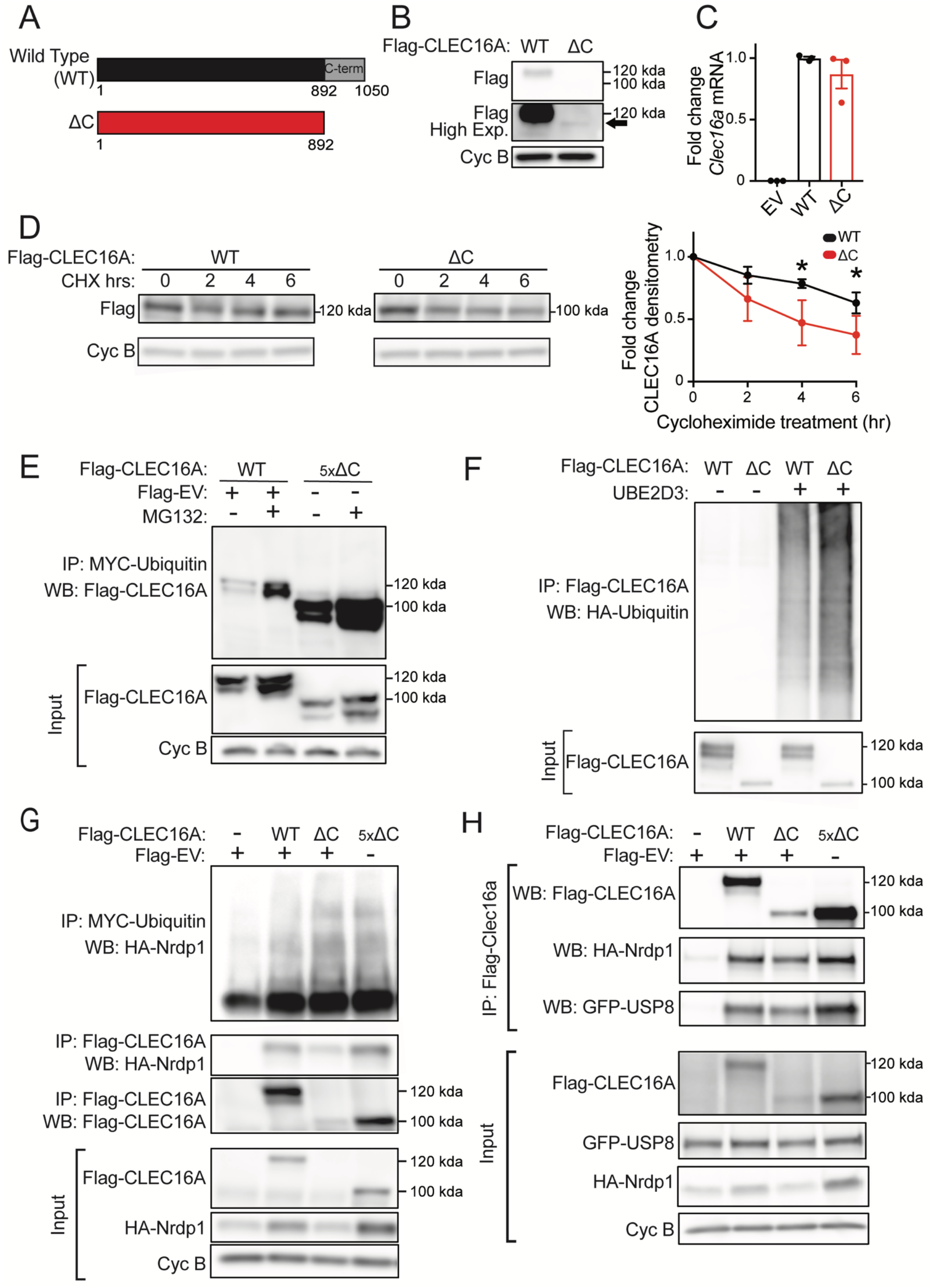
The CLEC16A C-terminal IDPR is required for mitophagy complex formation by maintaining CLEC16A stability. **(A)** Schematic of C-terminal Flag epitope-tagged constructs encoding wild type (WT) full-length CLEC16A or CLEC16A lacking the C-terminal IDPR (ΔC). **(B)** Representative Flag WB of WT and ΔC CLEC16A following transfection in 293T cells. Arrow indicates CLEC16A ΔC band. Cyclophilin B (Cyc B) serves as loading control. n=4/group. **(C)** Relative *Clec16a* mRNA levels (normalized to *PPIA* expression) following transfection of Flag-EV, WT CLEC16A, or ΔC CLEC16A plasmids into 293T cells. n=3/group. **(D)** Representative WB of Flag-CLEC16A levels (with densitometry of % change from basal levels) from 293T cells transfected with WT or ΔC CLEC16A following treatment with cycloheximide (CHX; 300µg/mL) for 0-6 hrs. Similar levels of Flag-CLEC16A protein between groups were achieved by transfection of 1µg of WT CLEC16A (with 4µg Flag-EV) or 5µg ΔC CLEC16A. n=3-4/group. **(E)** Representative WB of cell-based ubiquitination assay of overexpressed Flag-tagged CLEC16A WT or ΔC performed in HEK293T cells co-transfected with MYC-Ubiquitin. Cells were treated with DMSO or 10 μM MG132 for 12 hrs. Similar levels of Flag-CLEC16A protein between groups were achieved by transfection of 1x WT (1.5µg Flag-WT CLEC16A + 6µg Flag-EV) or 5x ΔC (7.5µg Flag-ΔC). n=3/group. **(F)** Representative WB of *in vitro* ubiquitination assay of recombinant CLEC16A-6xHis-Flag WT or ΔC following incubation with ATP, HA-Ubiquitin, E1, in the presence or absence of UbE2D3 at 37°C for 1 hour. n=4/group. **(G)** Representative WBs of cell-based assessment of binding, stabilization, and ubiquitination of overexpressed HA-tagged Nrdp1 by Flag-tagged CLEC16A WT or ΔC (or empty vector control) performed in HEK293T cells co-transfected with MYC-Ubiquitin. Conditions included Flag-EV (empty vector; 7.5µg), 1x Flag-CLEC16A WT (1.5µg Flag-CLEC16A WT + 6µg Flag-EV), and 1x or 5x Flag-CLEC16A ΔC (1.5µg Flag-CLEC16A ΔC + 6µg Flag-EV or 7.5 µg Flag-CLEC16A ΔC, respectively). n=3/group. **(H)** Representative WB following anti-Flag IP in HEK293T cells transfected with HA-Nrdp1, GFP-USP8 and Flag-EV, Flag-CLEC16A WT, or Flag-CLEC16A ΔC vectors. Conditions included Flag-EV (empty vector; 7.5µg), 1x Flag-CLEC16A WT (1.5µg Flag-CLEC16A WT + 6µg Flag-EV), and 1x or 5x Flag-CLEC16A ΔC (1.5µg Flag-CLEC16A ΔC + 6µg Flag-EV or 7.5µg Flag-CLEC16A ΔC, respectively). n=3/group. *p<0.05.

We next investigated the mechanism by which the CLEC16A C-terminal IDPR stabilizes CLEC16A. The stability of E3 ubiquitin ligases is commonly regulated by self-ubiquitination and subsequent degradation (46). Interestingly, terminal IDPRs in ubiquitin ligase complexes have been demonstrated to inhibit ubiquitin-chain assembly (47). We hypothesized that the CLEC16A C-terminal IDPR may moderate CLEC16A self-ubiquitination, thus protecting CLEC16A from degradation. Indeed, loss of the CLEC16A C terminus increased CLEC16A ubiquitination relative to WT CLEC16A when constructs were expressed at similar protein levels in HEK293T cells (Figure 4E). CLEC16A ΔC similarly had increased ubiquitination relative to WT CLEC16A following MG132 treatment (Figure 4E). To determine whether increased ubiquitination of CLEC16A ΔC was due to increased self-ubiquitination, we performed *in vitro* ubiquitination assays using recombinant CLEC16A. Indeed, loss of the CLEC16A C terminus increased CLEC16A self-ubiquitination relative to WT CLEC16A (Figure 4F). Together, these studies demonstrate that the CLEC16A C terminus protects CLEC16A from self-ubiquitination and degradation.

Based on the above findings, we questioned how loss of the CLEC16A C terminus and consequent CLEC16A destabilization would impact formation of the tripartite CLEC16A-Nrdp1-USP8 mitophagy complex (8). CLEC16A promotes formation of this complex by binding, ubiquitinating, and stabilizing the E3 ligase Nrdp1 (8). CLEC16A ΔC had impaired ability to bind and stabilize Nrdp1 relative to WT CLEC16A (Figure 4G). The impaired ability of CLEC16A ΔC to stabilize Nrdp1 was due to reduced levels of CLEC16A ΔC, as increasing CLEC16A ΔC protein levels back to that of WT CLEC16A restored the binding and stabilization of Nrdp1 (Figure 4G). Unexpectedly, CLEC16A C-terminal loss did not impair Nrdp1 ubiquitination despite reduced CLEC16A levels (Figure 4G). Loss of the CLEC16A C terminus also impaired formation of the mitophagy complex, as observed by reduced binding to USP8 and Nrdp1, which was again rescued by increasing CLEC16A ΔC protein levels back to that of WT CLEC16A levels (Figure 4H). Together, these studies indicate that the CLEC16A C terminus promotes mitophagy complex formation by stabilizing CLEC16A.

We next explored how the specific C-terminal deficiency in the human CLEC16A disease variant affected CLEC16A stability, ubiquitination, and mitophagy complex assembly. The CLEC16A disease variant had lower protein levels and was less stable than the full-length human CLEC16A isoform (Figures 5A-B). The CLEC16A disease variant was also more heavily ubiquitinated when protein levels were similar to that of the full-length isoform (Figure 5C), which was further increased following bafilomycin A_1_ treatment (Figure 5C; Figure S6B). The CLEC16A disease variant impaired formation of the mitophagy complex, demonstrated by reduced interaction between USP8-Nrdp1 (Figure 5D) as well as reduced stabilization of, and interaction with, Nrdp1 (Figure 5E). These results are similar to those of the CLEC16A ΔC mutant (Figure 4), which lead us to speculate that reduced stability of the CLEC16A disease variant is due to deficiency of the C-terminal IDPR.

**Figure 5.**
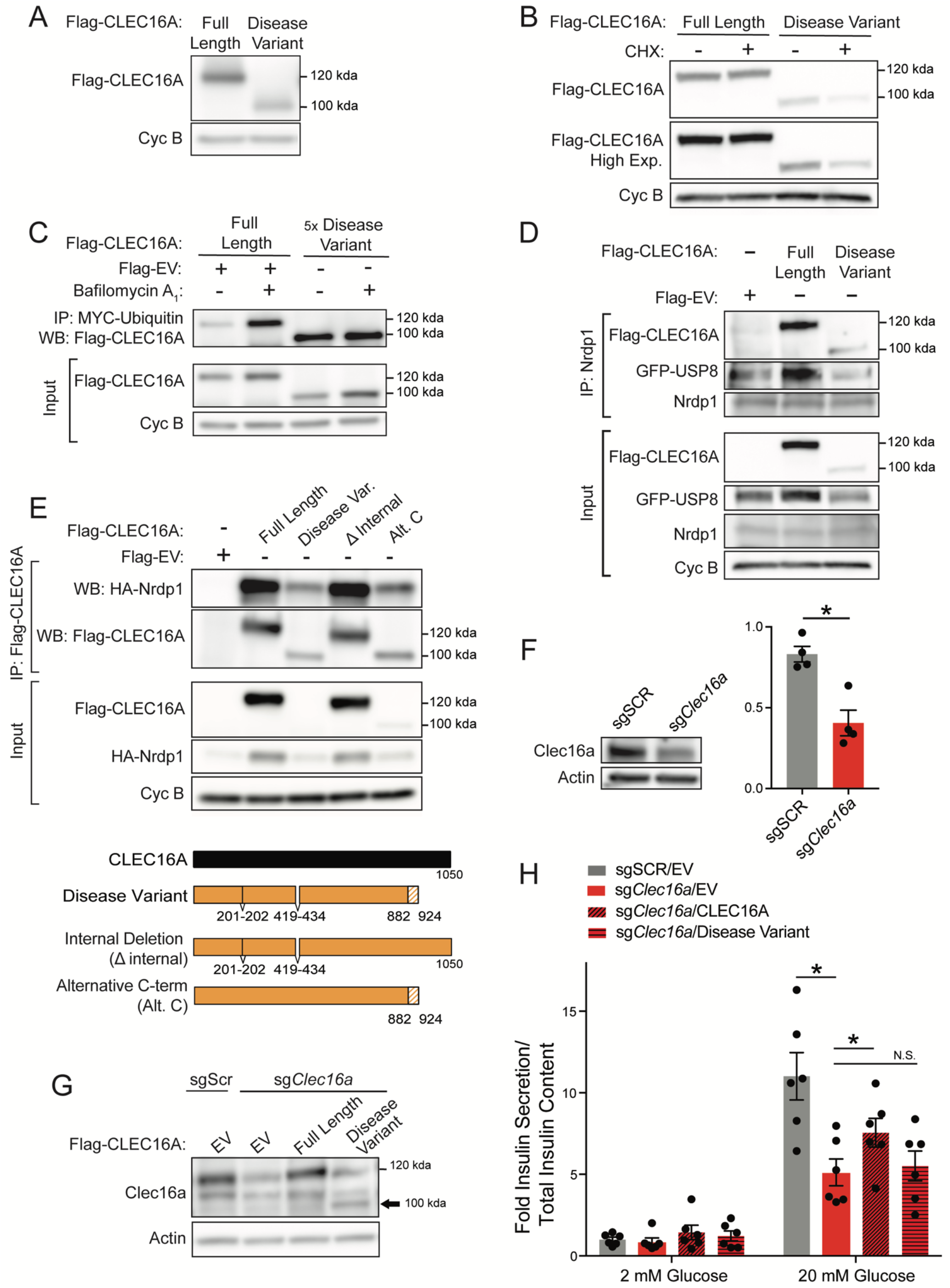
The human CLEC16A disease isoform is unstable and impairs mitophagy complex formation and β-cell function. **(A)** Representative Flag WB of full length human CLEC16A and the CLEC16A disease variant following transfection in 293T cells. n=3/group. **(B)** Representative WB of Flag-CLEC16A levels from 293T cells transfected with a plasmid encoding full-length human CLEC16A or the CLEC16A disease variant CLEC16A following treatment with cycloheximide (CHX; 300µg/mL) for 16 hours. n=3/group. **(C)** Representative WB of cell-based ubiquitination assay of overexpressed Flag-tagged CLEC16A or the CLEC16A disease variant performed in HEK293T cells co-transfected with MYC-Ubiquitin. Cells were treated with DMSO or 150 nM BafA for 12 hrs. Similar levels of Flag-CLEC16A protein between groups were achieved by transfection of the following plasmids encoding: 1x full-length CLEC16A (1.5µg Flag-CLEC16A + 6µg Flag-EV) or 5x CLEC16A disease variant (7.5µg Flag-CLEC16A disease variant). n=3/group. **(D)** Representative WB following endogenous Nrdp1 IP in HEK293T cells transfected with GFP-USP8 and Flag-EV, Flag-CLEC16A, or Flag-CLEC16A disease variant vectors. n=3/group. **(E)** Representative WBs of cell-based assessment of binding and stabilization of overexpressed HA-tagged Nrdp1 by Flag-tagged CLEC16A full length, CLEC16A disease variant, CLEC16A Δ internal, or CLEC16A alternative C-terminus (or empty vector control) performed in HEK293T cells co-transfected with MYC-Ubiquitin. n=3/group. **(F)** Representative CLEC16A WB of CLEC16A protein levels in scramble control (sgScr) or CLEC16A-deficient (sg*Clec16a*) Min6 cells generated by CRISPR-mediated gene editing (with densitometry). **(G)** Representative CLEC16A WB of Min6 control and CLEC16A-deficient cells from (F) following transfection with Flag-CLEC16A, Flag-CLEC16A disease variant, or empty vector control into CLEC16A-deficient Min6 cells. **(H)** Glucose stimulated insulin secretion (GSIS) assay performed in transfected Min6 cells from (G) 72 hours after transfection with CLEC16A (or empty vector control) constructs, following static incubation in 2 mM or 20 mM glucose. n=5/group. *p<0.05.

Beyond loss of the C terminus, the human CLEC16A disease variant also lacks two small internal regions (Figure 5E, lower panel). To clarify whether the disruption of internal or C-terminal regions drive the defects observed in the CLEC16A disease variant, we generated constructs bearing either the internal deletions (Δ internal) or the alternatively translated and truncated C terminus specific to the disease variant (alternative C terminus; Figure 5E, lower panel). While the CLEC16A Δ internal construct appeared to be functionally indistinguishable from full length CLEC16A, the human CLEC16A alternative C terminus construct had reduced protein levels and reduced ability to stabilize and bind Nrdp1 (Figure 5E). Overall, these results confirm that disrupting the C-terminal IDPR instigates the reduced stability and impaired mitophagy complex assembly found in the CLEC16A disease variant.

### The human CLEC16A disease variant is functionally defective in β-cells

Next, we wanted to determine if the mechanistic impairments observed in the human CLEC16A human disease variant would lead to β-cell dysfunction. To address the impact of the human CLEC16A disease variant and its disrupted C-terminal IDPR on β-cell function, we generated a Min6 β-cell line partially deficient for CLEC16A by CRISPR-Cas9 gene editing (Figure 5F). We then expressed either the full-length CLEC16A human isoform or the CLEC16A disease variant in CLEC16A-deficient Min6 cells (Figure 5G). As expected, CLEC16A deficiency reduced glucose-stimulated insulin secretion (GSIS) in Min6 cells (Figure 5H). Importantly, expression of the full length CLEC16A isoform, but not of the CLEC16A disease variant, improved GSIS in CLEC16A-deficient Min6 cells (Figure 5H). Thus, the human CLEC16A disease variant is functionally defective within β-cells, supporting a critical role for the CLEC16A C-terminal IDPR in β-cell function.

### Function of the CLEC16A C-terminal IDPR depends on its proline bias

After defining mechanistic and functional roles of the CLEC16A C-terminal IDPR, we questioned what features of the C-terminal IDPR mediate its effect on CLEC16A stability. While IDPRs lack secondary structure, IDPR function can be either dependent or independent of amino acid sequence order and composition (48; 49). IDPR functions that are sequence dependent include dependence on short linear motifs, post-translational modifications, charge, the distribution of charged amino acids, and amino acid biases, including specific behaviors of proline residues (49-54).

To first investigate whether CLEC16A C-terminal IDPR function depends on its amino acid sequence order, we generated two CLEC16A constructs with randomly shuffled C-terminal IDPR amino acids using the random shuffling tool, Shuffle Protein (55). We confirmed that these constructs did not eliminate predicted disorder within the C terminus (Figure 6A), nor did they introduce new protein domains, as determined using PROSITE and SMART domain analyses (56; 57). Levels of both CLEC16A C-terminal shuffled mutants were unchanged compared to WT CLEC16A (Figure 6B), indicating that CLEC16A stability is not dependent on the amino acid sequence order of its C-terminal IDPR.

**Figure 6.**
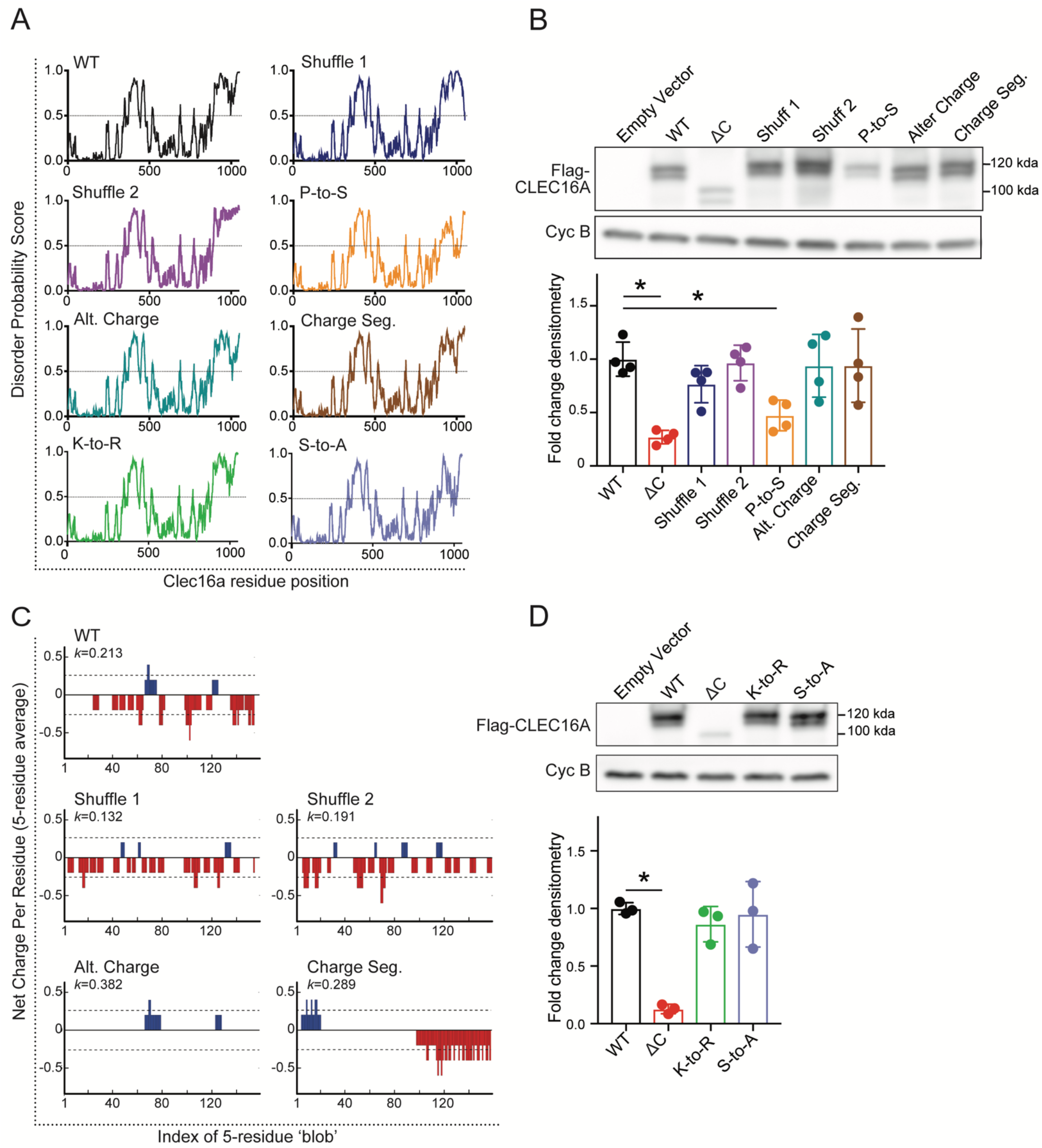
Proline bias within the CLEC16A C-terminal IDPR is necessary for CLEC16A stability. **(A)** Disorder propensity score for CLEC16A mutants as determined by IUPred2. **(B)** Representative Flag WB (with quantification by densitometry of all studies below) of Flag-CLEC16A WT, Flag-CLEC16A ΔC, as well as C-terminal IDPR mutants following transfection in 293T cells. n=4/group. C-terminal IDPR mutants include P-to-S (AA892-1050 proline to serine mutant), Alt. charge (alternate charge mutant), and Charge seg. (charge segregation mutant). **(C)** Net charge per-residue for listed CLEC16A mutants, calculated as an average net-charge per residue over a 5-residue ‘blob’. Kappa (*k*) scores indicates the degree to which charged residues are intermixed and were generated by CIDER (52). *p<0.05. **(D)** Representative Flag WB (with quantification by densitometry of all studies below) of Flag-CLEC16A WT, Flag-CLEC16A ΔC, as well as C-terminal pan-lysine to arginine (K-to-R) or pan-serine to alanine (S-to-A) mutants following transfection in HEK293T cells. n=3/group.

Next, we questioned whether the stabilizing function of the CLEC16A C-terminal IDPR depends on features of its amino acid composition. We generated several independent CLEC16A C-terminal mutants to assess whether disruption of certain IDPR features causes instability similarly to C-terminal truncation. We disrupted C-terminal proline bias with proline-to-serine mutagenesis, which conserves disorder but disrupts proline-specific interactions and behaviors (Figure 6A). We altered negative charge within the CLEC16A C terminus by aspartic acid to asparagine and glutamic acid to glutamine mutagenesis (Figure 6C). We disrupted the intermixed CLEC16A C-terminal charge distribution by manually segregating charged residues to alter the parameter kappa, which describes the intermixing of charges within a sequence (Figure 6C) (52). We also generated constructs with C-terminal regions resistant to post-translational modifications, including ubiquitination or phosphorylation, by lysine-to-arginine (K-to-R) and serine-to-alanine (S-to-A) mutagenesis, respectively. While CLEC16A stability did not depend on C-terminal charge, charge distribution, or post-translational modifications, it was dependent on the strong proline bias (Figures 6B,D). Proline residues are enriched in IDPRs, with unique properties due to their cyclical and rigid structure that disrupts secondary structure (32). Thus, the ability of the CLEC16A C-terminal IDPR to stabilize CLEC16A depends on an enrichment of proline residues.

## DISCUSSION

Here, we demonstrate that the CLEC16A C terminus is an IDPR that is required for CLEC16A function and glucose homeostasis. The CLEC16A C-terminal IDPR, which is disrupted in a human *CLEC16A* disease variant, is vital for mitophagy and insulin secretion by protecting CLEC16A from self-ubiquitination and degradation. The stabilizing role of the CLEC16A C-terminal IDPR relies on its proline bias, and not amino acid sequence order, charge, charge distribution, or post-translational modifications. Together, we use structural, mechanistic, and physiologic approaches to understand how an IDPR disrupted by a human disease variant regulates mitophagy and contributes to diabetes pathogenesis.

Our results provide new knowledge regarding the association between IDPRs and disease pathogenesis, which is poorly understood. While IDPRs comprise an estimated 44% of the human proteome, their biological roles have only recently begun to be explored (17; 58). IDPRs have proposed involvement with many human diseases, including cancer, cardiovascular disease, and neurodegeneration (17; 21; 24; 59). In some contexts, disease-causing mutations disrupt IDPRs, which suggest IDPRs have crucial biological functions that protect against disease (17; 21; 24; 59). Predicted IDPRs were enriched in a focused *in silico* analysis of 34 type 2 diabetes (T2D)-related proteins, yet the genetic, mechanistic, and physiological connections between IDPRs and the >400 variants associated with T2D are unknown (22). An IDPR was also biophysically validated in the β-cell transcription factor PDX1, but the functional contribution and genetic links of this IDPR to diabetes pathogenesis is unclear (35). To our knowledge, our study represents the first comprehensive structure-function characterization of an IDPR disrupted by a diabetes-associated human genetic variant.

We find that proline residues are required for CLEC16A C-terminal IDPR function, providing insight into how amino acid composition dictates IDPR function. Prolines are significantly enriched in IDPRs and tend to disrupt secondary structure due to their cyclical structure and lack of backbone amide hydrogen atoms, which prevents the formation of hydrogen bonds required for α-helices or β-sheets (32). Prolines have been implicated in controlling IDPR conformation, yet their role remains controversial and is likely context specific (31; 32; 60; 61). In some contexts, prolines are observed to promote expansion of IDPRs, with a positive correlation between proline number and IDPR hydrodynamic radii (60). Conversely, prolines are implicated in promoting compaction, as others found that mutating prolines within a short IDPR induced expansion (61). One mechanism by which proline residues can compact IDPRs is by interactions with proximal aromatic residues (31), yet the CLEC16A C terminus mostly lacks aromatic residues (Figures 1C,D). Future biophysical studies investigating how proline residues regulate the CLEC16A C-terminal IDPR conformation and behavior may clarify the role of proline bias in CLEC16A stability.

We also identify the first validated structural and functional domain within CLEC16A. Despite its known enzymatic activity as an E3 ubiquitin ligase, CLEC16A lacks a consensus E3 ligase domain (8). Our biophysical studies confirm the presence of a C-terminal IDPR, which stabilizes CLEC16A by attenuating its self-ubiquitination. Interestingly, the stabilizing role of the CLEC16A C-terminal IDPR opposes the well-known role for IDPRs to promote protein turnover through degradative post-translational modifications (45; 62; 63). We unexpectedly find the CLEC16A C-terminal IDPR enhances protein stability and reduces CLEC16A ubiquitination, indicating a novel function for this IDPR. The N-terminal IDPR of the E3 ligase RNF4 facilitates substrate ubiquitination in a manner dependent on IDPR compaction, which was controlled by charge segregation (64). However, the CLEC16A C-terminal IDPR is not required for ubiquitination of Nrdp1 (Figure 4G), nor does disruption of its charged residues affect CLEC16A stability (Figures 6A-C), together suggesting a distinct mechanism of action. Thus, our study of the CLEC16A C-terminal IDPR defines a surprising new role for IDPRs within E3 ligases. Functional exploration of other regions within CLEC16A, including a putative internal IDPR, will be of great interest in future work.

We describe molecular mechanisms by which a disease-associated CLEC16A variant impacts cellular function, which may provide opportunities for therapeutic interventions. Loss of the CLEC16A C-terminal IDPR destabilizes CLEC16A and impairs assembly of the tripartite mitophagy complex, which is overcome by increasing levels of the IDPR-deficient mutant (Figure 4H). Thus, pharmacological approaches to target the C-terminal IDPR to stabilize CLEC16A could prove protective against disease. Indeed, there has been growing interest and development in therapeutics targeting IDPRs (65). Interestingly, IDPRs can also retain functionality when expressed independently as truncated fragments (66-68). Exploring whether peptide-based therapeutics mimicking the structure of the CLEC16A C-terminal IDPR could improve CLEC16A stability and function would be of great value, not only to patients with diabetes, but also with other CLEC16A-related diseases.

## METHODS

### Protein Expression and Purification

The pMCSG7-Clec16a 892-1050-TEV-6xHis bacterial expression plasmid was transformed into Z-competent Rosetta2 cells (a gift from the Center for Structural Biology, University of Michigan), cultured in 10 mL Luria broth (LB) overnight, before transferring to 1L of LB and growing until optical density of 600 nm at 37°C. Bacteria was pelleted, washed, and resuspended in minimal medium (12 g/L K_2_HPO_4_, 9 g/L KH_2_PO_4_, 1 g/L ^15^NH_4_Cl, 2.5 g/L NaCl, 25 mg/L thiamine HCl, 4 g/L ^13^C-glucose, 0.5 g/L MgSO_4_, 0.1 g/L NaOH) supplemented with 1 mL of 100 mM CaCl_2_, and 1 mL of Metal Solution (0.3 g/65 mL FeSO_4_·7H_2_O, 0.2 g/65 mL ZnSO_4_·7H_2_O, 0.4 g/65 mL CoCl_4_·6H_2_O, 0.3 g/65 mL (NH_4_)_6_Mo_7_O_24_·4H_2_O, 0.3 g/65 mL CuSO_4_, 0.2 g/65 mL MnCl_4_·4H_2_O, 0.1 g/65 mL H_3_BO_3_), and ampicillin. Culture was induced overnight at 20°C with Isopropyl β-D-1-thiogalactopyranoside (IPTG). Bacteria were pelleted, subjected to freeze/thaw at -80°C, and sonicated in lysis buffer (1x PBS, 1% CHAPS, 10mM MgCl_2_, 2 µl benzonase (25 U/µl, Millipore), and protease inhibitor (Thermo Scientific)). Supernatant was purified using a nickel gravity column (Qiagen, Ni-NTA Agarose) followed by a high-salt wash (1xPBS, 1 M NaCl), and eluted in 1xPBS and 300 mM imidazole. Eluate was incubated with TEV protease overnight at 4°C in 1xPBS with 0.1% BME, and 6xHis tag was removed with a nickel column. Eluate was purified using Q anion exchange chromatography in the AKTAxpress (GE Healthcare), and fractions visualized on SDS-PAGE gel with Coomassie Brilliant Blue. Fractions containing the CLEC16A amino acids (AA) 892-1050 peptide were further purified using fast protein liquid size exclusion chromatography on the AKTA Purifier 10. Fractions were visualized on an SDS-PAGE gel and were dialyzed into the final NMR buffer (50 mM sodium cacodylate, 150 mM NaCl, pH 6.5 in ddH_2_O), followed by concentration.

For *in vitro* ubiquitination assays, recombinant CLEC16A and CLEC16A ΔC were generated following expression in 293T cells and purified with nickel-charged resin (Ni-NTA agarose; Qiagen) per the manufacturer’s protocols.

### NMR

Isotopically labeled ^13^C/^15^N CLEC16A C-terminal peptide (AA 892-1050) was purified using the methods described above and transferred into a buffer composed of 50mM sodium cacodylate pH 6.5, 150mM NaCl. Protein was supplemented with 10% D_2_O and loaded in a Shigemi NMR Tube (Wilmad-LabGlass, SP Scienceware). All NMR experiments were performed on an 18.8 T Bruker Ascend magnet equipped with Bruker NEO spectrometer operating at ^1^H frequencies of 800.25 MHz, and equipped with inverse TCI cryoprobe. All NMR spectra were collected at 298K.

Due to the limited lifetime of the sample, standard C’ detected NMR methods (29) were not suitable for chemical shift assignments for the CLEC16A C-terminal peptide (AA 892-1050). Initially a standard (H^α^-start) ^15^N-^13^C CON was collected using 1024(C’) x 128(N), spectral width of 11 × 36 ppm, and 16 scans. Some useful residue type information was obtained from the standard H(CC)CON from the Bruker pulse program library with matrix size of 1024(C’) × 128(N) × 48(C), spectral width of 8 × 36 × 80 ppm, and 16 scans with 25% non-uniform sampling, totaling an acquisition time of 19 hours. With the caveat of limited chemical shift dispersion in amide ^1^H dimension, we collected BEST-HNCO, BEST-HN(CA)CO, with a matrix size of 698(H_N_) × 64(N) × 128(C’), spectral width of 14 × 35 × 14 ppm, 8 and 16 scans respectively, and BEST-HNCACB and BEST-HN(CO)CACB, with a matrix size of 698(H_N_) × 64(N) × 128(C^α/β^), spectral width of 14 × 35 × 80 ppm, with 32 and 16 scans respectively, with 50% non-uniform sampling, totaling an acquisition of 30 hours for all 4 experiments. The backbone assignments were made by daisy-chain walking through C’ chemical shift assignments and aided by data from C^α/β^ chemical shifts as well. The aliphatic C^α/β^ chemical shifts also provided information on amino acid type owing to different chemical environment and types of side chains. Aliphatic protons chemical shifts from the H(CC)CON was also used to identify residue type, where data from BEST-HNCO was linked using C’ and N resonances.

Overall, spectra complexity and sequence repetitiveness made complete residue connection and assignment using this approach impossible. Combining fragmented data from C’- and H_N_-detect experiments described above, we assigned peaks to 16 of 159 total residues.

In lieu of assigning individual residues, carbon-detect amino acid selective (CAS) NMR experiments were performed, which leveraged the different chemical configuration of aliphatic side chains to assign aliphatic chemical shift resonances (35). Confirmation of assignments were made using the mapping of observed C’/N signals to those obtained from (H^α^-start) ^15^N-^13^C CON spectra. Carbon-detect amino-acid specific pulse sequences for serine, alanine, glycine, and threonine, were performed as described previously (35). Aliphatic C^α^ and C^β^ values were labeled for each peak on the CAS NMR spectra. All data were processed in Topspin 4.0.9 software and converted to Sparky format for data analysis.

### NMR-based secondary structure analyses

Secondary structure analyses of residue peaks with known amino acid type (identified using CAS-NMR) were performed by using secondary chemical shifts to evaluate the probability, for each residue, that the residue is in an α-helix or β-sheet (51). Data were generated by calculating the difference in the measured ^13^C-α and ^13^C-β chemical shift, from the expected chemical shift standards of that residue type in a random coil conformation, represented as dCα and dCβ. The difference between dCα and dCβ values (dCα-dCβ) was plotted, with values greater than 2 ppm or less than -2 ppm representing a residue likely in an α-helix or β-sheet, respectively.

### Animals

All animal studies were reviewed and approved by the University of Michigan Institutional Animal Care and Use Committee. Models used included *Clec16a*^curt/curt^ (38; 39) and mt-Keima mice (a gift from Dr. Toren Finkel, University of Pittsburgh (43)). *Clec16a*^curt^ mice (38; 39) were obtained from Jackson Labs and were maintained on a mixed 50% SWR/J and 50% CD1/ICR background. mt-Keima mice were maintained on a mixed SWR/J, CD1/ICR, and FVB/J background following intercrosses with *Clec16a*^curt/curt^ mice to generate experimental groups for mitophagy analysis. Mice were housed with a 12-h light/12-h dark cycle, and free access to food and water unless fasted for testing.

Mice were genotyped for the *Clec16a*^Curt^ allele with the following primers: Common forward primer: 5’-TGTCCCCATTACGCGTGCTAA-3’, and *Clec16a*^+^-specific reverse primer: 5’-TGTTGTCGGCTGGATTGGGA-3’, or *Clec16a*^Curt^-specific reverse primer: 5’-TGTTGTCGGCTGGATTGGTC-3’, which yield an 81 bp product. PCR was performed for 38 cycles (95°C for 45s, 58°C for 45s, 72°C for 1 min). Confirmation of the 4 bp deletion within exon 22 in *Clec16a*^Curt^ mice was performed following MnlI restriction digest of a PCR-amplified product of genomic DNA (forward primer: 5’-CCCCAAGGGTCTTACTGTCA-3’ and reverse primer: 5’-CATAGAAACGGAAAGGCAGGTGCTG-3’). *Clec16a*^+^-specific bands migrated at 137 and 47 bp, while a *Clec16a*^Curt^ mutant band migrated at 180 bp.

### Cell culture, cell line generation, treatments, and transfections

Primary mouse embryonic fibroblasts (MEFs) were isolated from *Clec16a*^Curt/Curt^ and *Clec16a*^+/+^ mice as described previously (69), and cultured as previously detailed (6; 70). 293T cells were cultured in DMEM supplemented with 10% Fetal-Plex (Gemini), 50 units/mL penicillin-streptomycin (Thermo Fisher Scientific), 1 mM sodium pyruvate (Thermo Fisher Scientific), and 142 μM β-mercaptoethanol (1 µl/100mL medium). Min6 cells were cultured as previously described (71). Pooled *Clec16a*-deficient Min6 cells were generated using the LentiCRISPRV2 one-vector system as described (72). *Clec16a* sgRNA sequences included (5’-CGGACATGTTTGGACGCTCA-3’, targeting exon 1) and (5’-ACAAAATCGGAACCTGCTCG-3’ targeting exon2) and were cloned into LentiCRISPRV2 as described (72). LentiCRISPRV2 was transfected into 293T cells along with the lentiviral packaging plasmids RRE, VSV-G, and REV (courtesy of Ling Qi, University of Michigan) using Lipofectamine 2000. Viral-containing media were collected on days 3 and 4 post-transfection, filtered in a 0.45 µm filter, and stored at -80°C until use. Min6 cells were transduced by culturing with viral media mixed 1:1 with normal Min6 media, supplemented with polybrene (5 µg/ml). Fresh viral stocks were added daily for two days, followed by two rounds of selection using puromycin (2 µg/mL) for 5 days. Min6 cells transduced with Clec16a sgRNA LentiCRISPR V2 (or non-targeting control sgRNA, courtesy of Ling Qi, University of Michigan) were screened via western blot. A pooled *Clec16a*-deficient Min6 cell population, and not an expansion of single cell *Clec16a*-deficient clones, was used to avoid bias from the variable insulin secretory profiles inherent to the generation of clonal β-cell lines.

The following compounds and concentrations were used as treatments for cells or isolated islets: dimethylsulfoxide (DMSO; Thermo Fisher Scientific), valinomycin (Millipore Sigma), MG132 (EMD Milipore), bafilomycin A_1_ (BafA, Cayman Chemicals), FCCP (Millipore Sigma), cycloheximide (EMD).

293T cells were transfected according to the manufacturer’s protocol using Lipofectamine 2000 (Thermo). Min6 cells were transfected using an Amaxa Nucleofector (Lonza) protocol G-016 according to the manufacturer’s protocol.

### Islet isolation and incubations

Islets were isolated from 12- to 15-week-old mice and subjected to static incubation to assess glucose-stimulated insulin secretion as previously described (71).

### Antibodies

CLEC16A-specific rabbit antisera (Cocalico) were generated as previously described, using either an internal peptide (AA 347-472) or a C-terminal peptide (AA 892-1050) (11). All other antibodies are listed in Table S1.

### Plasmids

Constructs for mammalian overexpression studies included, pFLAG-CMV-5a-Clec16a WT (6), pcDNA3.1 3x-HA-Nrdp1 (6), MYC-ubiquitin (71) (7), GFP-USP8 (Sino Biological; HG15979-ACG), and Human CLEC16A C-terminal Flag epitope-tagged full-length or alternatively spliced disease isoform vectors (Genscript; OHu18264D and OHu02258D). Constructs containing CLEC16A ΔC (1-892 only) or CLEC16A AA 892-1050 (C-terminal fragment only) were generated by PCR amplification of specific fragments from pFLAG-CMV-5a-Clec16a WT construct, subcloning of PCR amplified fragments into pCR-Blunt II-TOPO (Zero blunt TOPO cloning kit; ThermoFisher) for sequence validation by sequencing, and ligation of fragments into pFLAG-CMV-5a. Plasmids used to generate recombinant protein for in vitro ubiquitination assays included pFLAG-CMV-5a-Clec16a WT-6xHis (6) and pFLAG-CMV-5a-Clec16a ΔC-6xHis, generated by ligation of Clec16a ΔC (1-892 only) into pFLAG-CMV-5a-6xHis.

For the purpose of generating labeled recombinant CLEC16A C-terminal peptide fragment AA 892-1050 for NMR, the CLEC16A AA 892-1050 fragment was PCR amplified from pFLAG-CMV-5a-Clec16a WT with primers bearing ligation independent cloning sites (Fwd-5’-TACTTCCAATCCAATGCTTCTCCATCCCTGTCATCACC-3’, Rev-5’-TTATCCACTTCCAATGTTAGTGCTCTGTGGGTTCCG-3’. The product was then annealed with a linearized pMCSG7 bacterial plasmid (a gift from the Center for Structural Biology, University of Michigan) containing an N-terminal TEV-cleavable 6xHis tag and transformed into high efficiency DH5-α competent E. coli prior to confirmation by sequencing.

Plasmids bearing CLEC16A C-terminal IDPR mutation for mammalian overexpression studies were generated by gene synthesis into pBSK(+) (Biomatik) that were subsequently subcloned into pFLAG-CMV-5a, including pFLAG-CMV-5a-Clec16a_C-terminal_S-to-A (bearing serine to alanine mutagenesis of 31 serine residues in AA 892-1050), pFLAG-CMV-5a-Clec16a_C-terminal_Shuffle1, Shuffle 2, P-to-S, Alter Charge, and Charge Segregation (specific CLEC16A C-terminal mutant amino acid sequences are listed in Table S2). Randomly shuffled C-terminal mutants were generated to disrupt primary structure and any secondary structural features using the online Shuffle Protein tool, with PROSITE and SMART analyses used to ensure no domains were introduced during shuffling (55-57). The strong negative charge in the CLEC16A C terminus was disrupted by eliminating the negative charges with aspartic acid to asparagine and glutamic acid to glutamine mutagenesis, reducing the net charge per residue from -0.094 to 0.025, isoelectric point from 4.24 to 10.31, and retaining similarly structured amino acids. To disrupt the intermixed CLEC16A C terminus charge distribution, we manually segregated positively charged lysine and arginine residues towards the N terminus of this region, and negatively charged aspartic acid and glutamic acid toward the C terminus of this region, while leaving the remainder of the sequence intact. The impact was quantified with the parameter kappa, which describes charge intermixing (52). Kappa shifted from 0.213 in the WT C terminus to 0.294 in the charge-segregated construct. The pFLAG-CMV-5a-Clec16a_C-terminal_K-to-R plasmid was generated by site directed mutagenesis converting lysines 959 and 962 to arginines via QuikChange II Site Directed Mutagenesis kit, per the manufacturer’s instructions (Agilent). All plasmids were verified by sequencing.

To generate the CLEC16A mutant bearing only the alternative C terminus of the CLEC16A disease isoform, the C-terminal fragment of the CLEC16A disease isoform (OHu02258D; Genscript) was liberated by BamHI restriction digest, gel purified, and ligated in place of the C terminus of the full-length CLEC16A isoform construct (OHu18264D; Genscript). To generate the CLEC16A mutant bearing only the internal deletion of the CLEC16A disease isoform, the C-terminal fragment of the CLEC16A full-length isoform (OHu18264D), was liberated by BamHI restriction digest, gel purified, and ligated in place of the C terminus of the full length CLEC16A isoform construct (OHu02258D) leaving the internal deletion within the C-terminal disease isoform intact. Correct orientation of inserts was first assessed using HindIII/XhoI digest and then verified by sequencing.

### Western blotting, ubiquitination assays, immunoprecipitation

Western blots and ubiquitination assays were performed as previously described (6; 8). For immunoprecipitation (IP), cells were lysed in a protein lysis buffer consisting of 150 mM NaCl, 1% IGEPAL CA-630, and 50 mM Tris, pH 8.0, supplemented with protease and phosphatase inhibitors (Millipore), followed by shearing by passage through a 21-gauge needle while on ice. Lysates were clarified by centrifugation, then pre-cleared with protein G beads (Sigma). Protein lysates were incubated with either anti c-MYC agarose beads (Sigma A7470), anti c-MYC magnetic beads (Santa Cruz 500772), or Flag M2 Affinity Gel (Sigma A2220) overnight at 4°C. Beads were washed 3 times in lysis buffer, and conjugates were eluted in 2x Laemmli buffer (Sigma) at 70^°^C for 10 minutes prior to SDS-PAGE. IP for endogenous RNF41/Nrdp1 was performed as described (8).

### Flow cytometry

Flow cytometry of mt-Keima mouse islets was performed as previously described (73). In brief, isolated islets were cultured for one day, treated with valinomycin or DMSO, and dissociated into single cells using trypsin-EDTA (0.25%, Thermo). Cells were stained with DAPI (Thermo) and FluoZin-3 AM (Invitrogen; to detect zinc-enriched β-cells), and resuspended in phenol red-free media. Flow cytometric data was analyzed by FlowJo (BD). At least, 5,000 FluoZin-3-positive cells were analyzed from each islet preparation.

Mitophagy rates in MEFs was assessed using Mtphagy Dye (Dojindo) as previously described (73). MEFs were incubated with media supplemented with Mtphagy dye (100 nM) for 7.5 hours prior to flow cytometry analysis. Cells were stained with DAPI, then analyzed on the LSR Fortessa flow cytometer. Mtphagy signal was measured with excitation at 488 nm and emission at 710 nm. TMRE (Invitrogen) was excited with a 561-nm laser and emission was measured with a 574-nm filter to measure mitochondrial membrane potential. At least 19,000 live cells per sample were analyzed.

### β-cell mass, glucose, and insulin measurements

β-cell mass was calculated as previously described (71). IPGTT, ITT, and serum insulin measurements were obtained as described previously (74; 75). Pancreatic hormone content was analyzed via ELISA (ALPCO) following acid ethanol extraction as previously described (75). Static glucose-stimulated insulin secretion in Min6 cells was performed as previously described, and insulin was measured by ELISA (Mouse Ultrasensitive Insulin ELISA; ALPCO) (76).

### Respirometry

Oxygen consumption rates were measured in isolated islets using the XF96 Flux Analyzer (Seahorse Bioscience) according to the manufacturer’s protocol. Briefly, 20 dispersed islets were seeded per well of a Seahorse flux plate that was pre-treated with Cell-Tak (Corning), and islets were allowed to adhere overnight. Cells were then incubated for 1 hour at 37°C in atmospheric CO_2_ in pH 7.4 unbuffered RPMI1640 with 1.67 mM glucose (Seahorse bioscience). Islets were first stimulated with high glucose (16.7 mM) and then FCCP (5 µM) for oxygen consumption and extracellular acidification analyses. Oxygen consumption rates were normalized to total protein content, which was measured by MicroBCA (Pierce).

### RNA and DNA isolation, gene expression, and mtDNA analyses

RNA isolation and reverse transcription was performed as previously described (6; 77). DNA was isolated using the Qiagen DNeasy kit per the manufacturer’s protocol. Gene expression by quantitative RT-PCR and relative mtDNA:nuclear DNA ratios were measured as previously described (6; 77).

### Transmission electron microscopy

Isolated islets were washed 2x in PBS and fixed overnight in 2.5% glutaraldahyde at 4°C and pelleted in 2% agarose. Agarose plugs were washed in 0.1M cacodylate buffer (pH 7.2; CB), stained for 1 hour on ice with 1.5% K4Fe(CN)6 + 2% OsO4 in 0.1 M CB, washed in 0.1 M Na^2+^ acetate, pH 5.2, and en bloc stained in 2% uranyl acetate in 0.1 M Na^2+^ acetate, pH 5.2 for 1 hour. The plug was embedded and polymerized for 24hr at 70^°^C. Sections were cut with a diamond knife, and post-stained with 4% uranyl acetate and Reynolds’ lead citrate. Images were captured on a JEOL JEM-1400 transmission electron microscope.

### Statistics

Data are represented as mean values, with error bars denoting ± SEM. For all studies, statistical significance was determined using unpaired two-tailed student’s t-tests for single comparisons or two-way ANOVA (Prism GraphPad) for multiple comparisons testing. A post-hoc Sidak’s test was performed following ANOVA for samples reaching statistical significance. A 5% significance level was used for all statistical tests.

## Author Contributions

M.A.G. conceived, designed, and performed experiments, interpreted results, drafted and reviewed the manuscript. X.L., B.C., M.V., T.S., V.S., A.R., G.L.P., J.Z., and D.S. performed experiments, interpreted results, and reviewed the manuscript. D.J.K. and S.S. designed studies, interpreted results, and reviewed the manuscript. S.A.S. conceived and designed the studies, interpreted results, drafted, edited, and reviewed the manuscript.

## Acknowledgements

M.A.G. was supported by the NIH (F31-DK-122761, T32-GM-007315, T32-GM-008322). S.A.S was supported by the JDRF (CDA-2016-189, SRA-2018-539, COE-2019-861), the NIH (R01 DK108921, U01 DK127747), the Department of Veterans Affairs (I01 BX004444), the Brehm family, the Anthony family, and a Brehm T1D Pilot and Feasibility Grant from the Michigan Diabetes Research Center (P30-DK020572). G.L.P. was supported by the American Diabetes Association (19-PDF-063). S.S. was supported by the JDRF (CDA-2016-189) and NIH (R01 DK108921). The JDRF Career Development Award to S.A.S. is partly supported by the Danish Diabetes Academy and the Novo Nordisk Foundation. D.J.K. was supported by the NIH (GM131919). We acknowledge P. Blakely and J. Harrison in the Michigan Medicine Microscopy Core for advice and assistance with sample preparation and EM imaging. The generation of recombinant protein for biophysical studies reported in this publication was supported by the University of Michigan Center for Structural Biology (CSB). The CSB is grateful for support from the U-M Life Sciences Institute, the U-M Rogel Cancer Center, the U-M Medical School Endowment for Basic Sciences, and grants from the NIH. We thank the University of Michigan BioNMR Core for assistance performing, analyzing, and interpreting NMR studies. The University of Michigan BioNMR Core is supported by the U-M College of Literature, Sciences and Arts, Life Sciences Institute, College of Pharmacy and the Medical School along with the U-M Biosciences Initiative. We thank Drs. H. Popelka, E. Walker, P. Arvan, D. Fingar, and members of the Soleimanpour laboratory for helpful advice.

## REFERENCES

1. Hakonarson H, Grant SF, Bradfield JP, Marchand L, Kim CE, Glessner JT, Grabs R, Casalunovo T, Taback SP, Frackelton EC, Lawson ML, Robinson LJ, Skraban R, Lu Y, Chiavacci RM, Stanley CA, Kirsch SE, Rappaport EF, Orange JS, Monos DS, Devoto M, Qu HQ, Polychronakos C: A genome-wide association study identifies KIAA0350 as a type 1 diabetes gene. Nature 2007;448:591–594

2. Gingerich MA, Sidarala V, Soleimanpour SA: Clarifying the function of genes at the chromosome 16p13 locus in type 1 diabetes: CLEC16A and DEXI. Genes and immunity 2020;21:79–82

3. Fujimaki T, Kato K, Yokoi K, Oguri M, Yoshida T, Watanabe S, Metoki N, Yoshida H, Satoh K, Aoyagi Y, Nozawa Y, Kimura G, Yamada Y: Association of genetic variants in SEMA3F, CLEC16A, LAMA3, and PCSK2 with myocardial infarction in Japanese individuals. Atherosclerosis 2010;210:468–473

4. Hafler DA, Compston A, Sawcer S, Lander ES, Daly MJ, De Jager PL, de Bakker PI, Gabriel SB, Mirel DB, Ivinson AJ, Pericak-Vance MA, Gregory SG, Rioux JD, McCauley JL, Haines JL, Barcellos LF, Cree B, Oksenberg JR, Hauser SL: Risk alleles for multiple sclerosis identified by a genomewide study. N Engl J Med 2007;357:851–862

5. Yoshida T, Kato K, Yokoi K, Oguri M, Watanabe S, Metoki N, Yoshida H, Satoh K, Aoyagi Y, Nozawa Y, Yamada Y: Association of genetic variants with myocardial infarction in individuals with or without hypertension or diabetes mellitus. Int J Mol Med 2009;24:701–709

6. Soleimanpour SA, Gupta A, Bakay M, Ferrari AM, Groff DN, Fadista J, Spruce LA, Kushner JA, Groop L, Seeholzer SH, Kaufman BA, Hakonarson H, Stoffers DA: The diabetes susceptibility gene Clec16a regulates mitophagy. Cell 2014;157:1577–1590

7. Pearson GL, Gingerich MA, Walker EM, Biden TJ, Soleimanpour SA: A Selective Look at Autophagy in Pancreatic β-Cells. Diabetes 2021;70:1229–1241

8. Pearson G, Chai B, Vozheiko T, Liu X, Kandarpa M, Piper RC, Soleimanpour SA: Clec16a, Nrdp1, and USP8 Form a Ubiquitin-Dependent Tripartite Complex That Regulates beta-Cell Mitophagy. Diabetes 2018;67:265–277

9. Pearson G, Soleimanpour SA: A ubiquitin-dependent mitophagy complex maintains mitochondrial function and insulin secretion in beta cells. Autophagy 2018;14:1160–1161

10. Soleimanpour SA, Ferrari AM, Raum JC, Groff DN, Yang J, Kaufman BA, Stoffers DA: Diabetes Susceptibility Genes Pdx1 and Clec16a Function in a Pathway Regulating Mitophagy in β-Cells. Diabetes 2015;64:3475–3484

11. Sidarala V, Pearson GL, Parekh VS, Thompson B, Christen L, Gingerich MA, Zhu J, Stromer T, Ren J, Reck EC, Chai B, Corbett JA, Mandrup-Poulsen T, Satin LS, Soleimanpour SA: Mitophagy protects β cells from inflammatory damage in diabetes. JCI Insight 2020;5

12. Bradfield JP, Qu HQ, Wang K, Zhang H, Sleiman PM, Kim CE, Mentch FD, Qiu H, Glessner JT, Thomas KA, Frackelton EC, Chiavacci RM, Imielinski M, Monos DS, Pandey R, Bakay M, Grant SF, Polychronakos C, Hakonarson H: A genome-wide meta-analysis of six type 1 diabetes cohorts identifies multiple associated loci. PLoS genetics 2011;7:e1002293

13. Mero IL, Ban M, Lorentzen AR, Smestad C, Celius EG, Saether H, Saeedi H, Viken MK, Skinningsrud B, Undlien DE, Aarseth J, Myhr KM, Granum S, Spurkland A, Sawcer S, Compston A, Lie BA, Harbo HF: Exploring the CLEC16A gene reveals a MS-associated variant with correlation to the relative expression of CLEC16A isoforms in thymus. Genes and immunity 2011;12:191–198

14. Tian J, Wang Z, Mei S, Yang N, Yang Y, Ke J, Zhu Y, Gong Y, Zou D, Peng X, Wang X, Wan H, Zhong R, Chang J, Gong J, Han L, Miao X: CancerSplicingQTL: a database for genome-wide identification of splicing QTLs in human cancer. Nucleic acids research 2019;47:D909–d916

15. Buniello A, MacArthur JAL, Cerezo M, Harris LW, Hayhurst J, Malangone C, McMahon A, Morales J, Mountjoy E, Sollis E, Suveges D, Vrousgou O, Whetzel PL, Amode R, Guillen JA, Riat HS, Trevanion SJ, Hall P, Junkins H, Flicek P, Burdett T, Hindorff LA, Cunningham F, Parkinson H: The NHGRI-EBI GWAS Catalog of published genome-wide association studies, targeted arrays and summary statistics 2019. Nucleic acids research 2019;47:D1005–d1012

16. Liu Z, Huang Y: Advantages of proteins being disordered. Protein science:a publication of the Protein Society 2014;23:539–550

17. Babu MM: The contribution of intrinsically disordered regions to protein function, cellular complexity, and human disease. Biochemical Society transactions 2016;44:1185–1200

18. Fishbain S, Inobe T, Israeli E, Chavali S, Yu H, Kago G, Babu MM, Matouschek A: Sequence composition of disordered regions fine-tunes protein half-life. Nat Struct Mol Biol 2015;22:214–221

19. Chew LH, Lu S, Liu X, Li FK, Yu AY, Klionsky DJ, Dong MQ, Yip CK: Molecular interactions of the Saccharomyces cerevisiae Atg1 complex provide insights into assembly and regulatory mechanisms. Autophagy 2015;11:891–905

20. Wang B, Merillat SA, Vincent M, Huber AK, Basrur V, Mangelberger D, Zeng L, Elenitoba-Johnson K, Miller RA, Irani DN, Dlugosz AA, Schnell S, Scaglione KM, Paulson HL: Loss of the Ubiquitin-conjugating Enzyme UBE2W Results in Susceptibility to Early Postnatal Lethality and Defects in Skin, Immune, and Male Reproductive Systems. The Journal of biological chemistry 2016;291:3030–3042

21. Cheng Y, LeGall T, Oldfield CJ, Dunker AK, Uversky VN: Abundance of intrinsic disorder in protein associated with cardiovascular disease. Biochemistry 2006;45:10448–10460

22. Du Z, Uversky VN: A Comprehensive Survey of the Roles of Highly Disordered Proteins in Type 2 Diabetes. International journal of molecular sciences 2017;18

23. Kulkarni P, Uversky VN: Intrinsically Disordered Proteins in Chronic Diseases. Biomolecules 2019;9

24. Hegyi H, Buday L, Tompa P: Intrinsic structural disorder confers cellular viability on oncogenic fusion proteins. PLoS computational biology 2009;5:e1000552

25. Mei Y, Glover K, Su M, Sinha SC: Conformational flexibility of BECN1: Essential to its key role in autophagy and beyond. Protein science:a publication of the Protein Society 2016;25:1767–1785

26. Mistry J, Chuguransky S, Williams L, Qureshi M, Salazar GA, Sonnhammer ELL, Tosatto SCE, Paladin L, Raj S, Richardson LJ, Finn RD, Bateman A: Pfam: The protein families database in 2021. Nucleic acids research 2021;49:D412–d419

27. Erdős G, Dosztányi Z: Analyzing Protein Disorder with IUPred2A. Curr Protoc Bioinformatics 2020;70:e99

28. Hatos A, Hajdu-Soltész B, Monzon AM, Palopoli N, Álvarez L, Aykac-Fas B, Bassot C, Benítez GI, Bevilacqua M, Chasapi A, Chemes L, Davey NE, Davidović R, Dunker AK, Elofsson A, Gobeill J, Foutel NSG, Sudha G, Guharoy M, Horvath T, Iglesias V, Kajava AV, Kovacs OP, Lamb J, Lambrughi M, Lazar T, Leclercq JY, Leonardi E, Macedo-Ribeiro S, Macossay-Castillo M, Maiani E, Manso JA, Marino-Buslje C, Martínez-Pérez E, Mészáros B, Mičetić I, Minervini G, Murvai N, Necci M, Ouzounis CA, Pajkos M, Paladin L, Pancsa R, Papaleo E, Parisi G, Pasche E, Barbosa Pereira PJ, Promponas VJ, Pujols J, Quaglia F, Ruch P, Salvatore M, Schad E, Szabo B, Szaniszló T, Tamana S, Tantos A, Veljkovic N, Ventura S, Vranken W, Dosztányi Z, Tompa P, Tosatto SCE, Piovesan D: DisProt: intrinsic protein disorder annotation in 2020. Nucleic acids research 2020;48:D269–d276

29. Jones DT, Cozzetto D: DISOPRED3: precise disordered region predictions with annotated protein-binding activity. Bioinformatics (Oxford, England) 2015;31:857–863

30. Uversky VN: The intrinsic disorder alphabet. III. Dual personality of serine. Intrinsically disordered proteins 2015;3:e1027032

31. Mateos B, Conrad-Billroth C, Schiavina M, Beier A, Kontaxis G, Konrat R, Felli IC, Pierattelli R: The Ambivalent Role of Proline Residues in an Intrinsically Disordered Protein: From Disorder Promoters to Compaction Facilitators. Journal of molecular biology 2020;432:3093–3111

32. Theillet FX, Kalmar L, Tompa P, Han KH, Selenko P, Dunker AK, Daughdrill GW, Uversky VN: The alphabet of intrinsic disorder: I. Act like a Pro: On the abundance and roles of proline residues in intrinsically disordered proteins. Intrinsically disordered proteins 2013;1:e24360

33. Popelka H, Uversky VN, Klionsky DJ: Identification of Atg3 as an intrinsically disordered polypeptide yields insights into the molecular dynamics of autophagy-related proteins in yeast. Autophagy 2014;10:1093–1104

34. Tompa P: Intrinsically unstructured proteins. Trends Biochem Sci 2002;27:527–533

35. Sahu D, Bastidas M, Showalter SA: Generating NMR chemical shift assignments of intrinsically disordered proteins using carbon-detected NMR methods. Analytical biochemistry 2014;449:17–25

36. Marsh JA, Singh VK, Jia Z, Forman-Kay JD: Sensitivity of secondary structure propensities to sequence differences between alpha- and gamma-synuclein: implications for fibrillation. Protein science:a publication of the Protein Society 2006;15:2795–2804

37. Redmann V, Lamb CA, Hwang S, Orchard RC, Kim S, Razi M, Milam A, Park S, Yokoyama CC, Kambal A, Kreamalmeyer D, Bosch MK, Xiao M, Green K, Kim J, Pruett-Miller SM, Ornitz DM, Allen PM, Beatty WL, Schmidt RE, DiAntonio A, Tooze SA, Virgin HW: Clec16a is Critical for Autolysosome Function and Purkinje Cell Survival. Scientific reports 2016;6:23326

38. Harris BS, Fairfield HE, Reinholdt LG, Bergstrom DE, L.R.D: The first Clec16a mutant mouse exhibits defects in digits and tail. MGI Direct Data Submission. MGI 2013;J:190968

39. Harris B, Ward-Bailey P, Bergstrom D, Bronson R, Donahue L: Curvy tail: a new skeletal mutation that maps to Chromosome 16 MGI Direct Data Submission. MGI 2011;J:172931

40. Kaufman BA, Li C, Soleimanpour SA: Mitochondrial regulation of beta-cell function: maintaining the momentum for insulin release. Molecular aspects of medicine 2015;42:91–104

41. Suliman HB, Carraway MS, Tatro LG, Piantadosi CA: A new activating role for CO in cardiac mitochondrial biogenesis. J Cell Sci 2007;120:299–308

42. Piantadosi CA, Carraway MS, Babiker A, Suliman HB: Heme oxygenase-1 regulates cardiac mitochondrial biogenesis via Nrf2-mediated transcriptional control of nuclear respiratory factor-1. Circ Res 2008;103:1232–1240

43. Sun N, Malide D, Liu J, Rovira, II, Combs CA, Finkel T: A fluorescence-based imaging method to measure in vitro and in vivo mitophagy using mt-Keima. Nature protocols 2017;12:1576–1587

44. Darling AL, Uversky VN: Intrinsic Disorder and Posttranslational Modifications: The Darker Side of the Biological Dark Matter. Front Genet 2018;9:158

45. van der Lee R, Lang B, Kruse K, Gsponer J, Sánchez de Groot N, Huynen MA, Matouschek A, Fuxreiter M, Babu MM: Intrinsically disordered segments affect protein half-life in the cell and during evolution. Cell reports 2014;8:1832–1844

46. de Bie P, Ciechanover A: Ubiquitination of E3 ligases: self-regulation of the ubiquitin system via proteolytic and non-proteolytic mechanisms. Cell Death Differ 2011;18:1393–1402

47. Schumacher FR, Wilson G, Day CL: The N-terminal extension of UBE2E ubiquitin-conjugating enzymes limits chain assembly. Journal of molecular biology 2013;425:4099–4111

48. Keul ND, Oruganty K, Schaper Bergman ET, Beattie NR, McDonald WE, Kadirvelraj R, Gross ML, Phillips RS, Harvey SC, Wood ZA: The entropic force generated by intrinsically disordered segments tunes protein function. Nature 2018;563:584–588

49. Das RK, Ruff KM, Pappu RV: Relating sequence encoded information to form and function of intrinsically disordered proteins. Curr Opin Struct Biol 2015;32:102–112

50. Davey NE, Van Roey K, Weatheritt RJ, Toedt G, Uyar B, Altenberg B, Budd A, Diella F, Dinkel H, Gibson TJ: Attributes of short linear motifs. Mol Biosyst 2012;8:268–281

51. Bah A, Forman-Kay JD: Modulation of Intrinsically Disordered Protein Function by Post-translational Modifications. The Journal of biological chemistry 2016;291:6696–6705

52. Das RK, Pappu RV: Conformations of intrinsically disordered proteins are influenced by linear sequence distributions of oppositely charged residues. Proceedings of the National Academy of Sciences of the United States of America 2013;110:13392–13397

53. Firman T, Ghosh K: Sequence charge decoration dictates coil-globule transition in intrinsically disordered proteins. J Chem Phys 2018;148:123305

54. Martin EW, Holehouse AS, Grace CR, Hughes A, Pappu RV, Mittag T: Sequence Determinants of the Conformational Properties of an Intrinsically Disordered Protein Prior to and upon Multisite Phosphorylation. J Am Chem Soc 2016;138:15323–15335

55. Stothard P: The sequence manipulation suite: JavaScript programs for analyzing and formatting protein and DNA sequences. Biotechniques 2000;28:1102, 1104

56. Sigrist CJ, Cerutti L, Hulo N, Gattiker A, Falquet L, Pagni M, Bairoch A, Bucher P: PROSITE: a documented database using patterns and profiles as motif descriptors. Brief Bioinform 2002;3:265–274

57. Letunic I, Khedkar S, Bork P: SMART: recent updates, new developments and status in 2020. Nucleic acids research 2021;49:D458–d460

58. Oates ME, Romero P, Ishida T, Ghalwash M, Mizianty MJ, Xue B, Dosztanyi Z, Uversky VN, Obradovic Z, Kurgan L, Dunker AK, Gough J: D(2)P(2): database of disordered protein predictions. Nucleic acids research 2013;41:D508–516

59. Iakoucheva LM, Brown CJ, Lawson JD, Obradovic Z, Dunker AK: Intrinsic disorder in cell-signaling and cancer-associated proteins. Journal of molecular biology 2002;323:573–584

60. Marsh JA, Forman-Kay JD: Sequence determinants of compaction in intrinsically disordered proteins. Biophys J 2010;98:2383–2390

61. Glaves R, Baer M, Schreiner E, Stoll R, Marx D: Conformational dynamics of minimal elastin-like polypeptides: the role of proline revealed by molecular dynamics and nuclear magnetic resonance. Chemphyschem 2008;9:2759–2765

62. Dunker AK, Brown CJ, Lawson JD, Iakoucheva LM, Obradović Z: Intrinsic disorder and protein function. Biochemistry 2002;41:6573–6582

63. Pang CN, Hayen A, Wilkins MR: Surface accessibility of protein post-translational modifications. J Proteome Res 2007;6:1833–1845

64. Murphy P, Xu Y, Rouse SL, Jaffray EG, Plechanovová A, Matthews SJ, Carlos Penedo J, Hay RT: Functional 3D architecture in an intrinsically disordered E3 ligase domain facilitates ubiquitin transfer. Nature communications 2020;11:3807

65. Wichapong K, Silvestre-Roig C, Braster Q, Schumski A, Soehnlein O, Nicolaes GAF: Structure-based peptide design targeting intrinsically disordered proteins: Novel histone H4 and H2A peptidic inhibitors. Comput Struct Biotechnol J 2021;19:934–948

66. Sabari BR, Dall’Agnese A, Boija A, Klein IA, Coffey EL, Shrinivas K, Abraham BJ, Hannett NM, Zamudio AV, Manteiga JC, Li CH, Guo YE, Day DS, Schuijers J, Vasile E, Malik S, Hnisz D, Lee TI, Cisse, II, Roeder RG, Sharp PA, Chakraborty AK, Young RA: Coactivator condensation at super-enhancers links phase separation and gene control. Science (New York, NY) 2018;361

67. Kim MY, Na I, Kim JS, Son SH, Choi S, Lee SE, Kim JH, Jang K, Alterovitz G, Chen Y, van der Vaart A, Won HS, Uversky VN, Kim CG: Rational discovery of antimetastatic agents targeting the intrinsically disordered region of MBD2. Sci Adv 2019;5:eaav9810

68. Ambadipudi S, Zweckstetter M: Targeting intrinsically disordered proteins in rational drug discovery. Expert Opin Drug Discov 2016;11:65–77

69. Jozefczuk J, Drews K, Adjaye J: Preparation of mouse embryonic fibroblast cells suitable for culturing human embryonic and induced pluripotent stem cells. Journal of visualized experiments:JoVE 2012;

70. Seluanov A, Vaidya A, Gorbunova V: Establishing primary adult fibroblast cultures from rodents. Journal of visualized experiments:JoVE 2010;

71. Claiborn KC, Sachdeva MM, Cannon CE, Groff DN, Singer JD, Stoffers DA: Pcif1 modulates Pdx1 protein stability and pancreatic β cell function and survival in mice. The Journal of clinical investigation 2010;120:3713–3721

72. Sanjana NE, Shalem O, Zhang F: Improved vectors and genome-wide libraries for CRISPR screening. Nat Methods 2014;11:783–784

73. Sidarala V, Zhu J, Pearson GL, Reck EC, Kaufman BA, Soleimanpour SA: Mitofusins 1 and 2 collaborate to fuel pancreatic beta cell insulin release via regulation of both mitochondrial structure and DNA content. bioRxiv 2021:2021.2001.2010.426151

74. Sachdeva MM, Claiborn KC, Khoo C, Yang J, Groff DN, Mirmira RG, Stoffers DA: Pdx1 (MODY4) regulates pancreatic beta cell susceptibility to ER stress. Proceedings of the National Academy of Sciences of the United States of America 2009;106:19090–19095

75. Soleimanpour SA, Crutchlow MF, Ferrari AM, Raum JC, Groff DN, Rankin MM, Liu C, De León DD, Naji A, Kushner JA, Stoffers DA: Calcineurin signaling regulates human islet {beta}-cell survival. The Journal of biological chemistry 2010;285:40050–40059

76. Khoo C, Yang J, Rajpal G, Wang Y, Liu J, Arvan P, Stoffers DA: Endoplasmic reticulum oxidoreductin-1-like β (ERO1lβ) regulates susceptibility to endoplasmic reticulum stress and is induced by insulin flux in β-cells. Endocrinology 2011;152:2599–2608

77. Kolesar JE, Wang CY, Taguchi YV, Chou SH, Kaufman BA: Two-dimensional intact mitochondrial DNA agarose electrophoresis reveals the structural complexity of the mammalian mitochondrial genome. Nucleic acids research 2013;41:e58

