## Supplemental Figures and Tables for "The human disease gene *CLEC16A* encodes an intrinsically disordered protein region required for mitochondrial quality control"

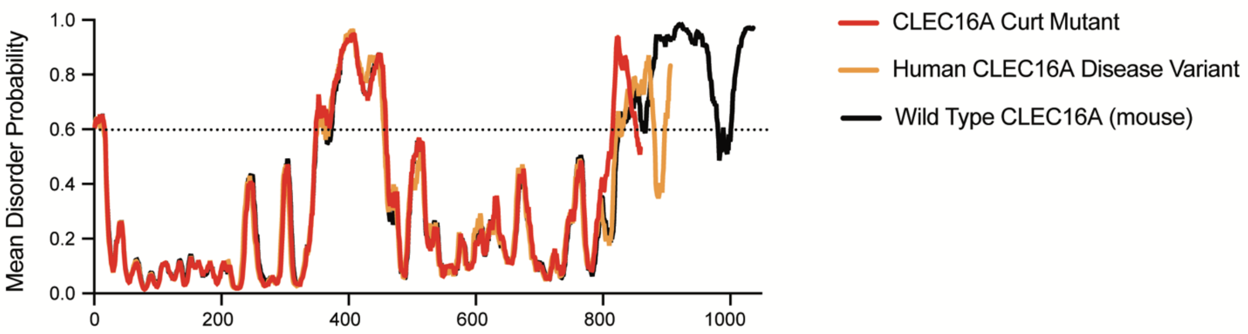


**Figure S1. The CLEC16A disease variant and CLEC16A-curt protein have similar predicted disorder profiles.** Mean disorder score from IUPred 2, Disprot VSL2B, and DISOPRED 3.1 of mouse full-length CLEC16A, the CLEC16A-curt protein, and the human CLEC16A disease variant. Two predicted disordered regions were identified by a probability threshold > 0.6.

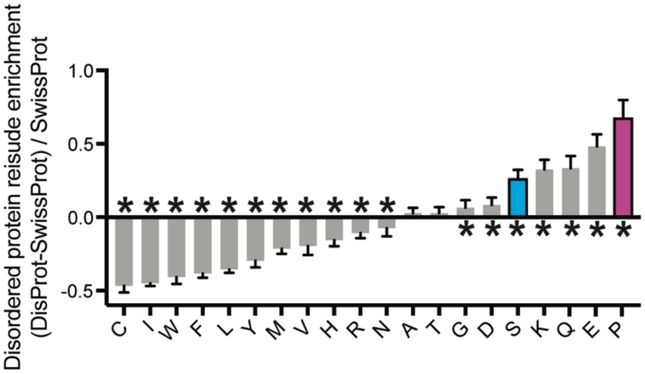


**Figure S2. Residue enrichment in disordered protein regions.** Residue composition bias of the verified disordered protein database DisProt generated with Composition Profiler. Compares residue enrichment in DisProt vs Swissprot51 database. Significantly enriched or depleted residues are indicated, * p<0.05.

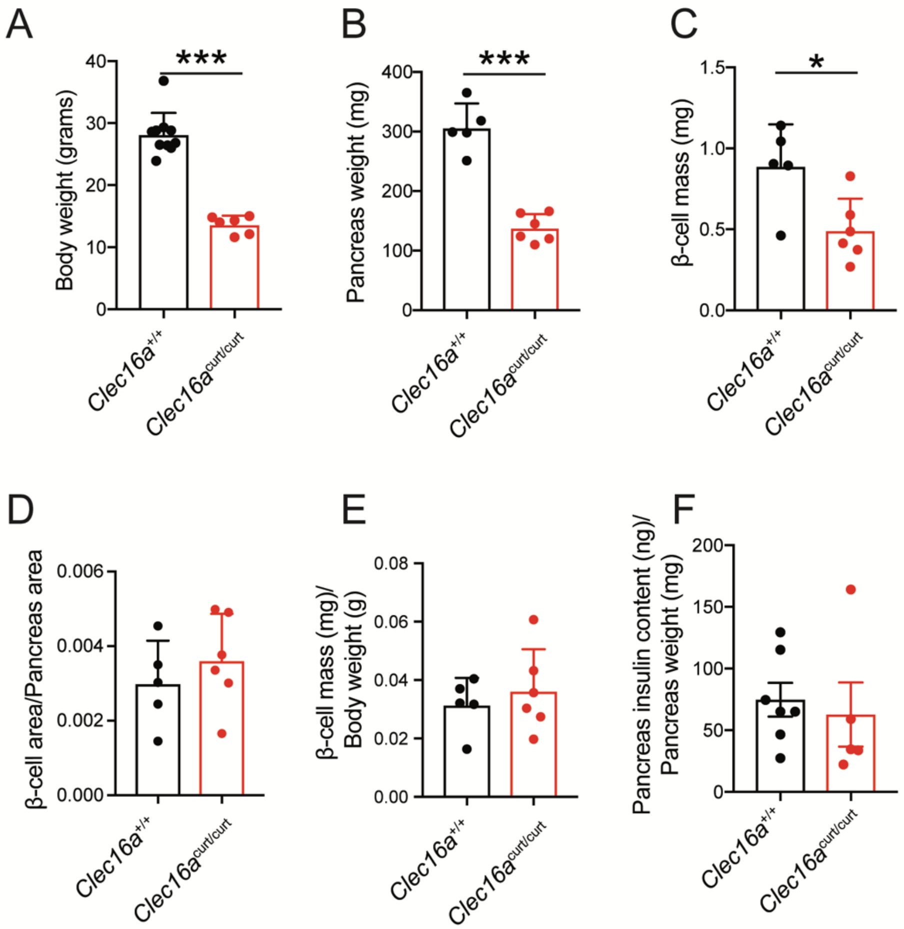


**Figure S3. CLEC16A C-terminal deficiency does not impair β-cell mass or insulin content relative to body or pancreas weight, respectively. (A)** Total body weight of 10-week old *Clec16a*$^{+/+}$ and *Clec16a*$^{curt/curt}$ mice. n=6-10/group. **(B)** Pancreas weight of 10-week old *Clec16a*$^{+/+}$ and *Clec16a*$^{curt/curt}$ mice. n=5-6/group. **(C)** β-cell mass of 10-week old *Clec16a*$^{+/+}$ and *Clec16a*$^{curt/curt}$ mice. n=5-6/group. **(D)** β-cell area normalized to pancreas area of 10-week old *Clec16a*$^{+/+}$ and *Clec16a*$^{curt/curt}$ mice. n=5-6/group. **(E)** β-cell mass normalized to body weight of 10-week old *Clec16a*$^{+/+}$ and *Clec16a*$^{curt/curt}$ mice. n=5-6/group. **(F)** Pancreatic insulin content normalized to pancreas weight of 10-week old *Clec16a*$^{+/+}$ and *Clec16a*$^{curt/curt}$ mice. n=5-6/group. *p<0.05, **p<0.01, ***p<0.001.

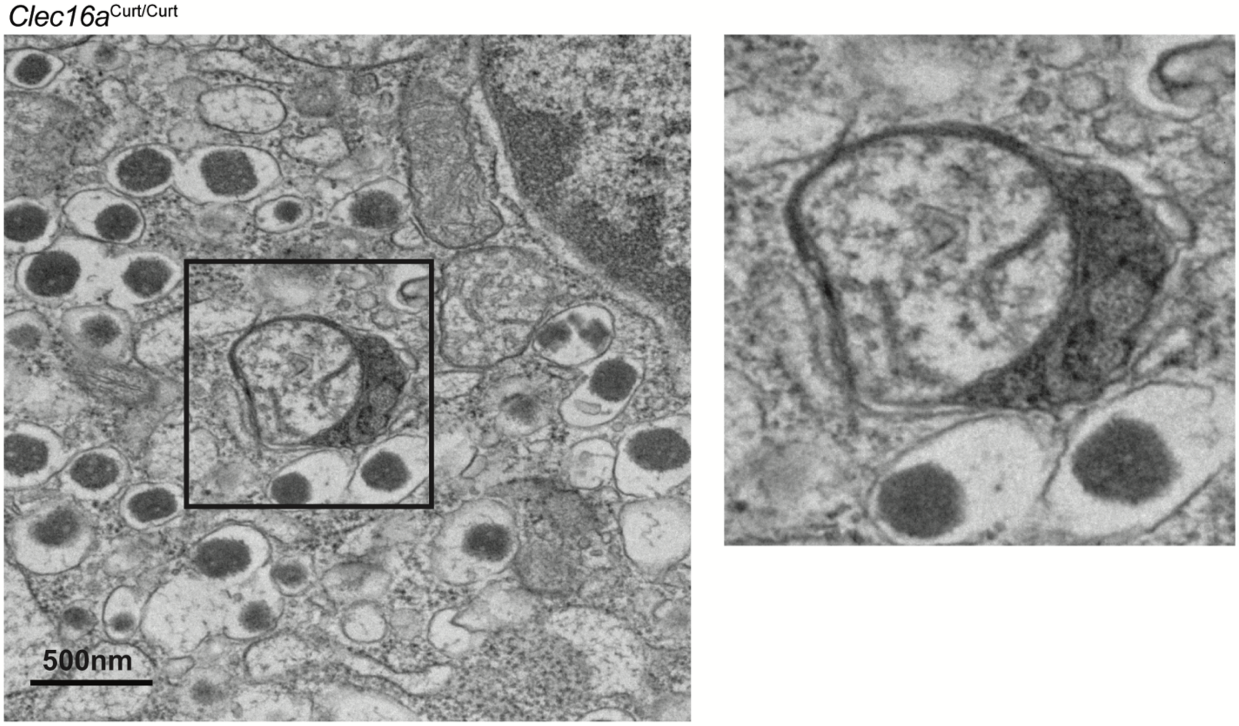


**Figure S4. *Clec16a*$^{curt/curt}$ β-cells demonstrate mitochondria in autophagosomal/lysosomal structures.** Representative transmission electron microscopy image of β-cell from 11-week-old *Clec16a*$^{curt/curt}$ islets from assessments in Figure 3A. Inset: focused areas of mitochondria surrounded by multiple membranes and undergoing mitophagy within a lysosomal structure.

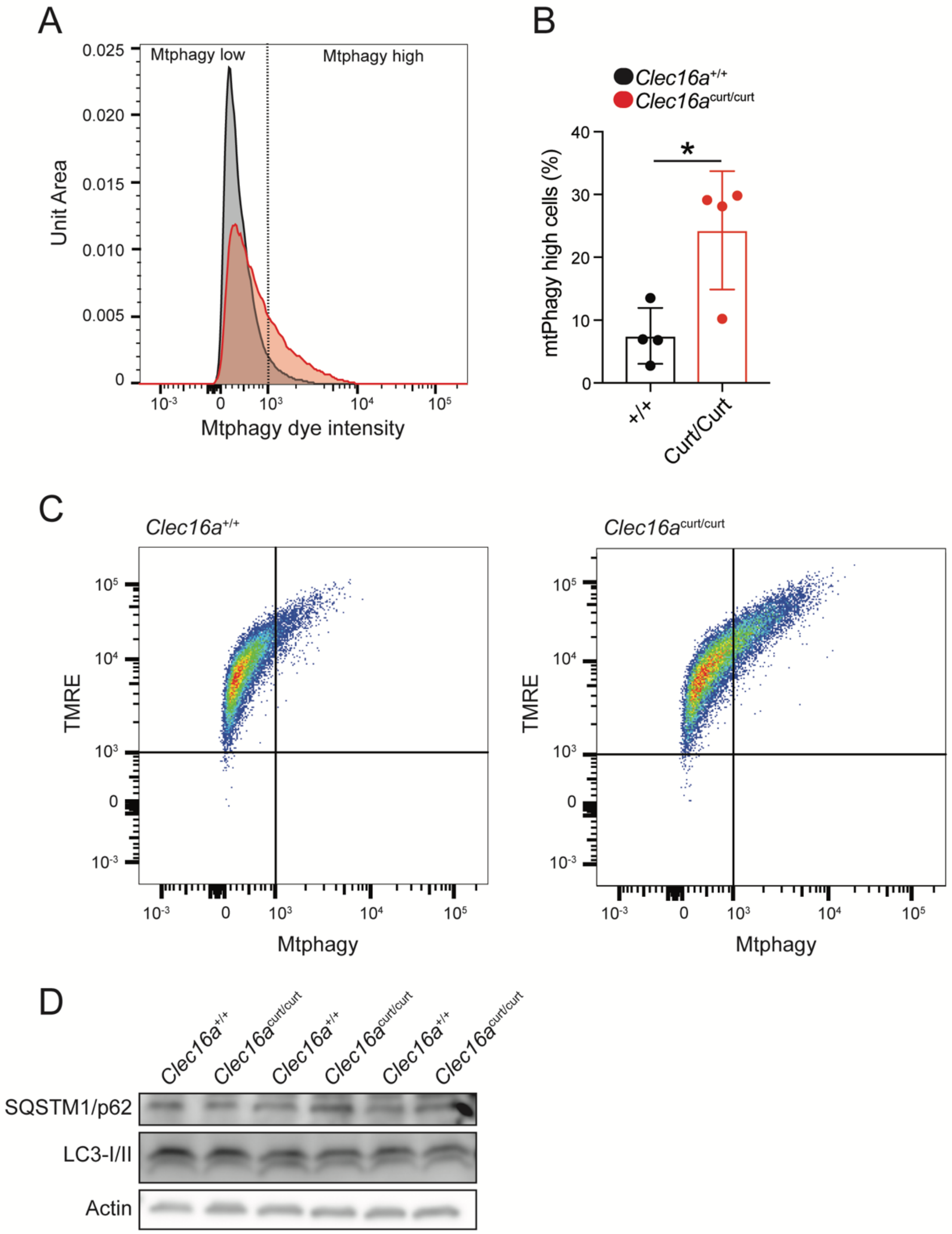


**Figure S5. Mitophagy is altered in *Clec16a*$^{curt}$ MEFS. (A)** Representative flow cytometry univariate histogram depicting Mtphagy dye intensity from primary MEFs derived from *Clec16a*$^{+/+}$ and *Clec16a*$^{curt/curt}$ mice. **(B)** Quantification of Mtphagy-high MEFs from *Clec16a*$^{+/+}$ and *Clec16a*$^{curt/curt}$ mice, generated using flow cytometry histograms as indicated in Figure S5A. n=4/group. **(C)** Representative flow cytometry scatter plot from *Clec16a*$^{+/+}$ and *Clec16a*$^{curt/curt}$ MEFs following staining with Mtphagy and TMRE dyes. **(D)** Western blot of autophagy markers SQSTM1/p62 and LC3 in *Clec16a*$^{+/+}$ and *Clec16a*$^{curt/curt}$ MEFs. n=3/group. *$p<0.05$

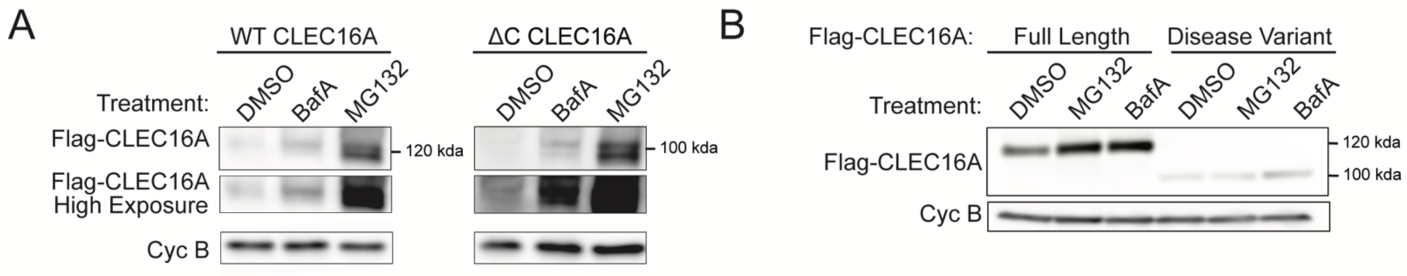


**Figure S6. Sensitivity of mouse CLEC16A and human CLEC16A to pharmacological inhibition of the proteasome or lysosome. (A)** Representative WB following transfection of plasmids encoding full-length mouse Flag-CLEC16A WT or Flag-CLEC16A ΔC in HEK293T cells treated for 12 h with 10 µM MG132, 150 nM bafilomycin $A_1$ (BafA), or DMSO. n=3/group. **(B)** Representative WB following transfection of plasmids encoding Flag-tagged full-length CLEC16A or CLEC16A disease variant in HEK293T cells treated for 12 h with 10 µM MG132, 150 nM bafilomycin $A_1$, or DMSO. n=3/group.

**Table S1**. Antibodies.

| **Antibody** | **Company** | **Catalog Number** |
|---|---|---|
| ACTB/β-Actin | ThermoFisher | MA5-15739 (BA3R) |
| Flag | Sigma | F1804 (clone M2) |
| GFP | Abcam | ab6673 |
| HA-Peroxidase | Roche | 12013819001 |
| Insulin | Dako | A0564 |
| LC3-I/II | Sigma | L8919 |
| MFN2 | Abcam | ab56889 (6A8) |
| P62 | Enzo | BML-PW9860 |
| Porin (VDAC1) | Abcam | ab14734 |
| PPIB/Cyclophilin B | ThermoFisher | PA1-027A |
| RNF41/Nrdp1 | Santa Cruz Biotechnology | sc-79996 (clone E-14) |
| SDHA | Abcam | ab14715 (2E3GC12FB2AE2) |
| Total OxPhos Rodent | Abcam | ab110413 |
| VCL/vinculin | Millipore | CP74-100UG |

**Table S2.** CLEC16A C-terminal IDPR mutagenesis sequences

| CLEC16A Construct | C terminus sequence (AA 892-1050) |
|---|---|
| WT | SPSLSSPSPPFASGSPGGSGSTSHCDSGGSSSAPSATQSPADAPTTPEQPQ PHLDQSVIGNEMDVNSKPSKNSSARSSEGETMHLSPSLLPAQQPTISLLYED TADTLSVESLTIVPPVDPHSLRALSGISQLPTLPAADTETPAEGAVNPEPAEPT EH |
| C-term S-to-A | APALAAPAPPFAAGAPGGAGATAHCDAGGAAAAPAATQAPADAPTTPEQPQ PHLDQAVIGNEMDVNAKPAKNAAARAAEGETMHLAPALLPAQQPTIALLYED TADTLAVEALTIVPPVDPHALRALAGIAQLPTLPAADTETPAEGAVNPEPAEPT EH |
| C-term K-to-R | SPSLSSPSPPFASGSPGGSGSTSHCDSGGSSSAPSATQSPADAPTTPEQPQ PHLDQSVIGNEMDVNSRPSRNSSARSSEGETMHLSPSLLPAQQPTISLLYED TADTLSVESLTIVPPVDPHSLRALSGISQLPTLPAADTETPAEGAVNPEPAEPT EH |
| C-term P-to-S | SSSLSSSSSSFASGSSGGSGSTSHCDSGGSSSASSATQSSADASTTSEQSQ SHLDQSVIGNEMDVNSKSSKNSSARSSEGETMHLSSSLLSAQQSTISLLYED TADTLSVESLTIVSSVDSHSLRALSGISQLSTLSAADTETSAEGAVNSESAEST EH |
| C-term Shuffle 1 | SSAPETLAHLALADLSESPPPGEAGTSGDSLLPLMNTLGTPEPISIQKSPECH PTAEVTKQLHEASPPGHDSPQPTNDASSGQAGSLHSFPITSPSGSDSTPPQ EVDPSTSPSSESPQNSSPEGTDSVLMPRSASAVTDQTNPAAGVPSYLVAIER LSS |
| C-term Shuffle 2 | SGAPASEEHQLPSGEGDAPLQSSPCLDNTKSAAASNLTQNSPSAGQIELADS DAPPPLHDTTLLRSEQEDVTSHTEPASLISPVSPRPQMAYHAEPPGIPTFTHQ SESVSSVPGKSSSNPLDGTSSVSEPSLPTSPIPSGVELSPPTSTPTMLSADA GG |
| C-term Alter Charge | SPSLSSPSPPFASGSPGGSGSTSHCNSGGSSSAPSATQSPANAPTTPQQP QPHLNQSVIGNQMNVNSKPSKNSSARSSQGQTMHLSPSLLPAQQPTISLLY QNTANTLSVQSLTIVPPVNPHSLRALSGISQLPTLPAANTQTPAQGAVNPQPA QPTQH |
| C-term Charge Segregation | SPSLSKSPSKPPFRASRGSPGGSGSTSHCSGGSSSAPSATQSPAAPTTPQP QPHLQSVIGNMVNSPSNSSASSGTMHLSPSLLPAQQPTISLL YTATLESVSL ETIEVPPVDPHEDSLDALESGDISQELPTDLPDAADTETPAEGADVNPEPAEP TEH |
